# Gluten-free Fish? Marine Carnivores Cobia (*Rachycentron canadum*) and European Sea Bass (*Dicentrarchus labrax*) Have Different Tolerances to Dietary Wheat Gluten

**DOI:** 10.1101/669655

**Authors:** Mary E.M. Larkin, Aaron M. Watson, Allen R. Place

## Abstract

In developing more sustainable fishmeal-free diets for a broad range of fish species, a “one-size-fits-all” approach should not be presumed. The production of more ecologically sustainable aquaculture diets has increased the incorporation of plant-based protein sources such as wheat gluten. Here we show that wheat gluten at even less than 4% inclusion in a compound feed has a negative impact on growth and survivorship in juvenile cobia (*Rachycentron canadum*). In addition, plasma factors capable of binding wheat gluten were detected in the plasma of cobia fed diets containing this ingredient but not in wild cobia with no exposure to dietary wheat gluten. Furthermore, there is evidence that supplementary taurine partially mitigates the deleterious effects provoked by wheat gluten. Based on these results, we propose that wheat gluten should be added with caution to aquaculture diets intended for juvenile cobia and potentially other marine carnivores. After observing that dietary wheat gluten can cause deleterious effects in cobia, we sought to evaluate a possible effect in European sea bass (*Dicentrarchus labrax*), another large, carnivorous, marine species. There were no major effects in terms of growth rate, plasma biochemical parameters, or detectable induction of plasma IgM, IgT, or factors capable of binding gliadin in response to 4% dietary wheat gluten. However, plasma levels of taurine doubled and there were considerable changes to the intestinal microbiome. There was increased diversity of predominant taxonomic orders in the pyloric caeca, anterior, middle, and posterior intestinal sections of fish consuming wheat gluten. Despite these measurable changes, the data suggest that dietary inclusion of 4% wheat gluten is well tolerated by European sea bass in feed formulations. Together these findings underscore the need to evaluate tolerance to ingredients in aquaculture formulations on a species by species basis.

## Introduction

Reducing fishmeal inclusion in favor of more sustainable and cost-effective protein sources is a high priority for the aquaculture industry (1). Aquaculture diets that incorporate low-fat, high-protein concentrates derived from plants such as soy or wheat (including processed wheat gluten) have been described for many species (2)(3). Feed formulations often incorporate wheat gluten as a protein source and for binding and adding a desirable chewy texture to pellets. Wheat flour is processed to remove soluble fibers and starches, and the vital wheat gluten product that remains contains two fractions: soluble gliadins and insoluble glutenins (4)(5). Vital wheat gluten, marketed as a dry powder, regains its elastic properties when rehydrated (6). In addition to being a low-cost protein source, it is useful for the binding together of feed ingredients into granules and pellets. Gliadins are known to trigger an immune response in susceptible people, such as those with celiac disease (7–9).

When plant proteins replace fishmeal, essential amino acids (e.g. lysine, methionine, threonine), as well as various vitamins and minerals, often need to be supplemented to counteract deficiencies in the protein source (10–12). One such necessary nutritional constituent found in fishmeal but is absent in plants is taurine. Some fish species are inadequate synthesizers of taurine and require dietary intake (13). Previous work by our laboratory showed that cobia is one such species, and it is essential to add taurine to a plant-based feed formulation intended for their consumption (14). Findings presented in this work suggest that taurine may help to mitigate some of the deleterious effects prompted by dietary wheat gluten in cobia.

The original intent of the cobia study described here was to test various plant-based diets formulated from available and cost-effective ingredients that had been previously found to be highly digestible by cobia and to be effective fishmeal replacements in rainbow trout (*Oncorhyncus mykiss*) and other species (14). Commercial feed formulations tend to vary based on the availability and cost of ingredients, with different batches potentially containing significantly different proportions and quality of ingredients, or even different ingredients altogether. The assessment of multiple components and combinations is necessary in the development of optimized fishmeal replacement diets. It is also necessary to identify which components might need to be supplemented for particular species when plant proteins replace fishmeal in a formulation.

In a previous study, when a plant protein-based diet lacking taurine was fed to cobia, poor growth and palatability were observed. However, when taurine was added to a similar plant protein-based diet, growth performance was as good if not better than a fishmeal-based commercial diet (14). It is likely that taurine supplementation greatly contributed to the improved performance of the diet. However, two of the plant protein sources were also replaced: barley meal (10.4%) and wheat gluten (8.2%) in exchange for solvent-extracted soybean meal (12.1%) and wheat flour (22.7%). The barley meal was replaced because of poor digestibility (< 60%), and the wheat gluten was replaced with wheat flour as part of a slight reformulation unrelated to specific concerns regarding wheat gluten. In the current study, which was not originally designed to evaluate wheat gluten as an ingredient, we prepared plant-based diets for cobia with varying concentrations of taurine (0 to 5%) at a fixed vital wheat gluten level of 2.2%. Wheat flour in the formulations also contributed a small amount of wheat gluten. Dietary wheat gluten at less than 4% inclusion appeared to be poorly tolerated, causing poor growth and even mortalities in juvenile cobia. We also detected the presence of a plasma factor(s) capable of binding gliadin that was present only in cobia exposed to dietary wheat gluten. Our laboratory wished to expand on this study to further delineate the impacts of wheat gluten and taurine but were unable to procure another cohort of cobia. We decided to investigate the impact of dietary wheat gluten in European sea bass (*Dicentrarchus labrax*), another marine carnivore.

European sea bass are a globally important aquaculture fish, with approximately 60,000 tons of fish produced each year as part of a greater than 300 million dollar market. They can be found on restaurant menus as branzino or branzini (15). A study in European sea bass suggests that they perform well on a diet containing wheat gluten (∼25% of formulation) when fishmeal is also included in the formulation (16). Another study included 20% gluten in an entirely plant-based formulation and noted differences in growth rates between the fish consuming that diet as compared to a fishmeal-based diet (17). It is important to note than in the dietary formulations for the study, the added wheat flour contributes a small amount of gluten. However, results from cobia suggest that wheat flour does not elicit the same negative effects as the processed vital wheat gluten. Being uncertain as to the taurine requirements of European sea bass, we supplemented the diets in the study with 1.5% taurine to ensure that taurine deficiency would not be a confounding factor in evaluating the effects of wheat gluten in European sea bass.

In the current study, we sought to determine if the inclusion of 4% wheat gluten into the diet of European sea bass impacted overall growth, health and immune status, and the intestinal microbiome. This entailed tracking growth rates, assaying plasma parameters and tissue weights, detecting adaptive immune factors IgM and IgT, and evaluating differences in the intestinal microbiome. European sea bass, like other teleost fish, have both innate and adaptive immunity. European sea bass and other teleosts have three types of immunoglobulins: IgM, IgT, and IgD (18, 19). The functions of the first two types have been well described. The role of IgM in fish is similar to its role in other organisms. It is the principal immunoglobulin found in teleost plasma and though its primary role is in systemic immunity, a role in mucosal immunity has also been described. IgT is an integral part of mucosal immunity and functions similarly to IgA in mammals. Like IgM, it is detectable in plasma, though levels tend to be lower (20).

Regarding the gut microbiome, it is known that predominant intestinal microbes in fish by diet, including with the addition of plant-based protein sources such as soybean meal (21–23). A study of European sea bass fed combined fish- and plant-based protein sources reported the main gut genera to be *Lactobacillus*, *Pseudomonas*, *Vibrio*, and *Burkholderia* (24). In this study, we wished to characterize both the microbiome of European sea bass fed a plant-based diet and the contribution of wheat gluten to the microbial landscape.

European sea bass fed the 0 and 4% wheat gluten diets had similar growth rates and levels of common plasma parameter, and neither group had detectable induction of plasma IgM, IgT, or factors capable of binding gliadin. Notable differences between the groups included twice the plasma levels of taurine in the fish fed 4% wheat gluten and considerable differences in the predominant taxonomic orders of the intestinal microbiome. Despite these measurable changes, the data suggest that dietary inclusion of 4% wheat gluten is well tolerated by European sea bass in an aquaculture feed formulation.

## Materials and Methods

### Fish System Maintenance and Care

#### Cobia

This study was carried out in accordance with the guidelines of the International Animal Care and Use Committee of the University of Maryland Medical School (IACUC protocol #0610015). Approximately 500 juvenile (∼2 g) cobia were obtained from the Virginia Agricultural Experiment Station, Virginia Tech, Hampton, VA, USA, for the first and second trials and approximately 500 juveniles (∼2 g) were obtained from the University of Miami in Miami, FL, USA, for the third trial. Juveniles were housed at the Institute of Marine and Environmental Technology’s Aquaculture Research Center in Baltimore, MD. Fish for the first and second trials (PP1-PP4) were maintained on a fishmeal diet until they reached an average weight of ∼10 g (Trial 1) or ∼120 g (Trial 2), at which point 18 fish were stocked into each of 12 identical tanks and randomly assigned one of the four experimental diets using three replicate tanks per dietary treatment. Six 340-liter tanks connected to bubble-bead and biological filtration as well as protein skimmers constituted the recirculating systems. Four replicate systems were occupied simultaneously during trials with the photoperiod maintained at 14 hours light, 10 hours dark throughout the trials. The first trial was conducted for 8 weeks, with tank weights recorded and feeding rates adjusted weekly to 5% bw day^-1^. The second trial commencing with ∼120 g fish was conducted for 8 weeks, with tank weights recorded weekly and feeding rates adjusted from 3.5% bw day^-1^ to 2.5% bw day^-1^, with a bi-weekly 0.25% bw day^-1^ reduction throughout the trial.

The third trial (EPP3 diet) was initiated with ∼18 g average weight individuals, with 12 fish stocked per tank. The third trial was conducted for 12 weeks, with tank weights recorded and feeding rates adjusted weekly to 5% bw day^-1^ for the first 6 weeks, reduced to 3.5 % from 6 weeks through 10 weeks, and 3.0% for the final 2 weeks of the trial as feed conversion ratio gradually increased. Fish received the daily ration by hand over the course of 4 feedings.

#### European sea bass

This study was carried out in accordance with the guidelines of the International Animal Care and Use Committee of the University of Maryland Medical School (IACUC protocol #0616014). European sea bass were obtained from the laboratory of Yonathan Zohar, PhD, and maintained in the Aquaculture Research Center at the Institute of Marine and Environmental Technology. Starting at ∼25 g of body weight, fish were fed diets containing either 0 or 4% wheat gluten (Zeigler Bros., Gardners, PA, USA), starting with 75 fish per diet. Fish were maintained on these diets for 6 months, at which time the average weight was 250 g. Fish were fed 3.5% of their body weight per day over 3-4 feedings.

Temperature was maintained at 27 degrees C and salinity was 25 ppt. Fish were divided by diet and housed in one of 2 eight-foot diameter, four cubic meter, recirculating systems sharing mechanical and bio-filtration as well as life support systems. The recirculating system has a filtration system which including protein skimming, ozonation, mechanical filtration in the form of bubble-bead filters, and biological filtration. Water samples were tested 2-3 times per week and analyzed by the National Aquarium water quality lab at IMET. Water quality was not significantly different between systems utilized (ANOVA, p>0.05) during the study and overall parameters were: dissolved oxygen, 5.69 ± 1.62 mg/L; temperature, 26.85 ± 1.77 degrees C; pH (4,500-H^+^), 7.61 ± 0.27; total ammonia nitrogen (4,500-NH_3_), 0.06 ± 0.06 mg/L; nitrite (4,500-NO_2_^-^), 0.12 ± 0.08 mg/L; nitrate (4,500-NO_3_^-^), 49.28 ± 8.87 mg/L, alkalinity (2, 320), 95.77 ± 23.11 meq/L; and salinity (2, 510) 24.91 ± 1.65 ppt.

### Diet preparation

#### Cobia

Formulations of the five plant protein (PP) diets used in the feed trials are shown in S1 Table: Feed formulations for the cobia dietary study. For all diets, ingredients were ground using an air-swept pulverizer (Model 18H, Jacobsen, Minneapolis, MN) to a particle size of < 200 μm. All ingredients for PP1, PP2, PP3, and PP4 were mixed prior to extrusion, while EPP3 was top-coated with the oil ingredient after extrusion. Pellets were prepared with a twin-screw cooking extruder (DNDL-44, Buhler AG, Uzwil, Switzerland) with an 18-second exposure to 127 °C in the extruder barrel. Pressure at the diet head was approximately 26 bar, and a die head temperature of 71 °C was used. The pellets were dried for approximately 15 minutes to a final exit air temperature of 102 °C using a pulse bed drier (Buhler AG, Uzwil, Switzerland) followed by a 30 min cooling period to product temperature less than 25 °C. Final moisture levels were less than 10% for each diet. Diets were stored in plastic lined paper bags at room temperature and were fed within six months of manufacture. Portions of each diet were analyzed by New Jersey Feed Labs, Inc. (Trenton, NJ, USA) for proximate composition (S2 Table: Proximate composition and measured taurine values of the cobia diets). Calculations of feed gluten content are based on wheat flour containing 8% gluten plus any added vital wheat gluten (25)

#### European sea bass

The formulations for the 0 and 4% wheat gluten diets are shown in S3 Table: Feed formulations for the wheat gluten European sea bass dietary study (Zeigler Bros., Gardners, PA, USA). The proximate composition of the feeds in shown in S4 Table: Proximate composition and measure taurine values of the European sea bass diets. The analysis was performed by New Jersey Feed Laboratory, Inc. (Ewing Township, NJ, USA).

### Blood and Tissue Sampling and Analysis

#### Cobia

At the conclusion of the first trial, two individuals from each tank were sacrificed for intestinal analysis. Portions of the anterior intestine were preserved in 4% paraformaldehyde and dehydrated from 70% to 90% EtOH in 10% increments over eight hours. Dehydrated samples were sent to AML Laboratories (Baltimore, MD) for sectioning, mounting, and H&E staining. Slides were analyzed for pathologies and abnormalities with the aid of the acknowledged pathologist, Dr. Renate Reimschuessel, VMD, PhD, (FDA in Laurel, MD). Gall bladders were removed and bile was extracted and stored at −20 °C prior to bile salt analysis. Total bile salts were assayed with 3 α-hydroxysteroid dehydrogenase (26). Blood samples were taken from the caudal vein with heparinized needles, plasma was separated by centrifugation (16,000 x g for 20 min), and total plasma protein was quantified after a 1:600 dilution utilizing a Micro BCA™ Protein Assay Kit (ThermoFisher Scientific, Waltham, MA, USA).

At the conclusion of the second trial, two fish from each tank (total of six per dietary treatment), were randomly selected for sampling. Fish were anesthetized with Tricaine methanosulfonate (MS-222, 70 mg L^-1^, Finquel, Redmond, WA, USA), blood samples were taken from the caudal vein with heparinized needles, after which fish were euthanized with MS-222 (150 mg L-^1^) and gall bladders were removed for bile analysis as in Trial 1. Liver and fillet samples were also taken for histology. Blood plasma was separated by centrifugation (16,000 x g for 20 min at 4 °C) and plasma osmolality measured in triplicate (10 μl) on a Vapro™ Model 5520 vapor pressure osmometer (Wescor, Logan, UT, USA). Plasma samples from three fish per dietary treatment were sent to the Pathology and Laboratory Medicine Services department at the University of California at Los Angeles for constituent analysis. Remaining plasma, fillet, and liver samples were frozen and stored at −80 °C and portions of each were lyophilized to constant weight for water and taurine content analysis. Triplicate samples of each liver (∼10 mg), fillet (∼50 mg), plasma (∼10 μL), and diet (∼50 mg) sample were used for taurine extractions based on Chaimbault et al., with samples homogenized in cold 70% EtOH, sonicated for 20 min, dried, and resuspended in 1 ml H_2_O prior to injection into the LC-MS (27).

#### European sea bass

At the conclusion of the 6-month trial, food was withheld for 24 hours and 10-12 fish from each diet were sedated with 25 mg/L MS-222 (Syndel, Ferndale, WA, USA) buffered with 50 mg/L sodium bicarbonate (Sigma-Aldrich, St. Louis, MO, USA) and exsanguinated via the caudal vein to collect blood for plasma analysis. Following blood collection, the spinal cord was severed, and tissues were harvested for analysis.

Approximately 1 ml blood from the caudal vein was put into a tube containing 20 µL of 1000 units/ml heparin and gently inverted to mix. Samples were centrifuged at 2000 x g for 15 minutes at 4 degrees C and the plasma fraction retained. This processing was sufficient for plasma chemistry and immunoblotting. In preparation for taurine analysis by HPLC, 10 µL plasma was mixed with 90 µL (1:10) 70% ethanol containing 0.154 mM D-norleucine. The solution was vortexed and centrifuged at 2000 x g for 5 min. 50 µL supernatant was retained and dried down at 70 degrees C overnight.

#### Plasma Analysis

Plasma analysis was performed by Jill Arnold of the National Aquarium in Baltimore, MD. Samples were processed using standard procedures for biochemistry analytes (S5 Table: Common plasma analytes measured) using the ChemWell-T analyzer (CataChem, Oxford, CT, USA). Calibration and quality control materials were used per manufacturer’s instructions (Catacal and Catatrol control level 1 and 2, CataChem, Oxford, CT, USA). Osmolality was measured using the Wescor Osmometer (Wescor, Inc., Logan, UT, USA) after calibration with two levels of standards (290 and 1000 mmol/kg, OPTIMOLE, ELITechGroup Biomedical Systems, Logan, UT, USA).

#### Plasma taurine analysis by HPLC

Dry samples were resuspended in 300 µL 0.1 N HCl and filtered through 0.45 micron filters (EMD Millipore, Billerica, MA, USA). 5 µL of the filtered extracts was derivatized according to the AccQTag™ Ultra Derivitization Kit protocol (Waters Corporation, Milford, MA, USA). Amino acids were analyzed using an Agilent 1260 Infinity High Performance Liquid Chromatography System equipped with ChemStation (Agilent Technologies, Santa Clara, CA, USA) by injecting 5 µL of the derivatization mix onto an AccQTag™ Amino Acid Analysis C18 (Waters, Milford, MA, USA) 4.0 um, 3.9 x 150 mm column heated to 37 °C. Amino acids were eluted at 1.0 mL min^-^1 flow with a mix of 10-fold diluted AccQTag™ Ultra Eluent (C) (Waters Corporation, Milford, MA, USA), ultra-pure water (A) and acetonitrile (B) according to the following gradient: Initial, 98.0% C/2.0% B; 2.0 min, 97.5% C/2.5% B; 25.0 min, 95.0% C/5.0% B; 30.5 min, 94.9% C/5.1% B; 33.0 min, 91.0% C/9.0% B; 38 min, 40.0% A/60.0% B; 43 min, 98.0% C/2.0% B. Derivatized amino acids were detected at 260 nm using a photo diode array detector. Signals were referenced to AABA (alpha-Aminobutyric acid), D-norleucine, and standard hydrolysate amino acids.

#### SDS-PAGE and western blotting

A solution of gliadin (Sigma, St. Louis, MO, USA) was made to an original concentration of 2 mg/ml and subsequently diluted 2-fold to 1 mg/ml, 0.5 mg/ml, 0.25 mg/ml, and 0.125 mg/ml, all in Laemmli sample buffer. All samples were heated to 95 °C for 3 min and centrifuged at 10,000 x g for 1 min prior to electrophoresis. Recombinant *Amphidinium carterae* eIF4A-1A (obtained from Grant Jones at IMET, 7/28/14) was diluted into Laemmli sample buffer to a concentration of 57.5 ng/µl. 15 µl of a 1:1 volume ratio of gliadin and eIF4A was loaded into each lane of a Novex NuPAGE 4-12% Bis-Tris gel and electrophoresed in a Bolt^®^ Mini Gel Tank at 165 V for 1 h with MOPS SDS running buffer (Life Technologies, Frederick, MD, USA).

PVDF membrane was activated by a brief dip in 100% methanol and equilibrated for 5 min in Novex™ NuPAGE^®^ transfer buffer. The gel was electroblotted onto a prepared PVDF in a Bolt Mini Blot Module at 30 V for 1 h in Novex™ NuPAGE^®^ transfer buffer (Life Technologies, Frederick, MD, USA). Following transfer, the membrane was washed in ddH_2_0 for 5 min. Duplicate transferred lanes were divided and designated as “No Plasma Block” or “With Plasma Block” for subsequent incubation procedures.

For the “No Plasma Block” sections, the membrane was incubated at room T for 1 h followed by overnight at 4 °C with 5 ml 5% nonfat milk in TBS-T. For cobia, the “With Plasma Block” membrane was incubated at room T for 1 h followed by overnight at 4 °C with either mixed plasma from a total of 7 fish from Trial 2 fed diets containing 3.2-3.6% gluten (PP1-PP4) or mixed plasma from two wild-caught cobia diluted 1:10 into TBS-T containing 5% nonfat milk. For European sea bass, the “0 Wheat Gluten Plasma” and “4% Wheat Gluten Plasma” membranes were incubated at room T for 1 h followed by overnight at 4 °C with mixed plasma from a total of 3 fish fed diets containing either 0 wheat gluten or 4% wheat gluten, respectively, diluted 1:12 into TBS-T containing 5% nonfat milk. The following day, blots were washed 4 times for 10 min each time with TBS-T. Both blots were incubated with polyclonal anti-gliadin antibody (Biorbyt, Cambridge, UK) and rabbit anti-*A. carterae* eIF4E-1A (GenScript, Piscataway, NJ, USA) diluted 1:500 and 1:2000, respectively, into TBS-T containing 5% nonfat milk at room temperature for 1 h. The blots were again washed 4 times for 10 min each time with TBS-T. Both blots were incubated with goat anti-rabbit IgG-HRP conjugate (Bio-Rad, Hercules, CA, USA) diluted 1:2500 into TBS-T containing 5% nonfat milk at room temperature for 1 h followed by four 10-min washes with TBS-T. The HRP signal from bound antibody was visualized using Clarity™ Western ECL Substrate (Bio-Rad, Hercules, CA, USA). Imaging was performed in a Flourchem™, and the AlphaView program was used to analyze densitometry (ProteinSimple, San Jose, CA, USA).

#### IgM and IgT immunoblotting in European sea bass

Plasma was diluted 1:40 in 1x SDS-PAGE sample buffer. Samples were heated for 3 minutes at 95 degrees C and centrifuged for one minute at 10,000 x g. 13 µL of each sample was electrophoresed on a 4%–12% Bis-Tris protein gel (NuPAGE Novex, (ThermoFisher Scientific, Waltham, MA, USA) for 35 minutes at 200 V using MOPS buffer in a PowerPac™ HC Power Supply (Bio-Rad, Hercules, CA, USA. Proteins were transferred to a PVDF membrane for 14 minutes on the high molecular weight setting (25 V) in the Trans-Blot® Turbo™ Transfer System (Bio-Rad, Hercules, CA, USA). Immunoblotting was performed in the iBind™ Western System (ThermoFisher Scientific, Waltham, MA, USA).

For IgT detection, an anti-European sea bass IgT polyclonal antibody (rabbit IgG “RAIgT1,” kindly provided by Giuseppe Scapigliati, Tuscia University, Italy) was used at a dilution of 1:1000 as the primary antibody, and goat anti-rabbit IgG H&L HRP conjugate at a dilution of 1:2000 (Bio-Rad, Hercules, CA, USA) was used as the secondary antibody. For IgM detection, the Magic™ anti-European sea bass IgM monoclonal antibody (mouse IgG) (Creative Diagnostics, Shirley, NY, USA) was used at a dilution of 1:1000 as the primary antibody, and goat anti-mouse IgG H&L HRP conjugate (Bio-Rad, Hercules, CA, USA) was used as the secondary antibody at a dilution of 1:2000. A chemiluminescent signal was generated with addition of Clarity™ Western ECL substrate and imaged in a ChemiDoc™ Touch Imaging System (Bio-Rad, Hercules, CA, USA). Image Lab software (Version 5.2.1, Bio-Rad, Hercules, CA, USA) was used to visualize immunoblots and analyze protein molecular weights.

### Microbiome analysis in European sea bass

#### DNA Extraction

For sampling of tank water, 1 L of water was filtered through a pore size of 0.2 microns. For the feed, 0.127 g (3 pellets) of each diet was used. For intestinal samples (pyloric caeca, anterior intestine, mid-intestine, and posterior intestine), ∼0.25 g tissue was collected. DNA extraction was performed using the Qiagen DNeasy® Powerlyzer® Powersoil® kit (Qiagen, Germantown, MD, USA). Samples were manually processed through the beat beating step 6 m/s for 30 sec, repeated once (FastPrep®-24, MP Biomedicals, Santa Ana, CA, USA) and centrifugation procedure for two minutes at 10,000 x g (Eppendorf 5415 D, Sigma-Aldrich, St. Louis, MO, USA). In the case of the water analysis, the filter was treated as the sample for processing. Following Step 5 in the manufacturer’s protocol, DNA extracted was completed in a Qiacube® (Qiagen, Germantown, MD, USA) according to manufacturer’s instructions for DNA extraction including the optional PCR inhibitor removal. DNA was stored at −20 degrees C.

#### PCR and Gel Electrophoresis

PCR was performed to confirm the presence of amplifiable DNA for 16s rRNA sequencing. One µL of extracted DNA was used in a 25 µL reaction volume with Promega™ PCR Master Mix (Thermo Fisher Scientific, Waltham, MA) and amplified in a DNA Engine Dyad (MJ Research, Quebec, Canada) using primers 16S_27F 5’ AGAGTTTGATCMTGGCTCAG 3’ and 16S_1492R 5’ TACGGYTACCTTGTTACGACTT 3’ with the following reaction conditions: 95 degrees C for 5 min, 92 degrees C for 30 sec, 50 degrees C for 2 min, 72 degrees C for 1 min, 30 sec, cycle to step 2 for 39 more times, incubate at 72 degrees C for 5 minutes, hold at 4 degrees C. Agarose gel electrophoresis was performed on each PCR product to detect the presence of a 1465 bp DNA band spanning bacterial rRNA variable regions 1-9. Samples were electrophoresed in a 1% agarose gel containing 0.5 μg/mL ethidium bromide at 150V for 40 minutes and imaged in a ChemiDoc™ Touch Imaging System (Bio-Rad, Hercules, CA, USA).

#### Sequencing

Sequencing was performed at the BioAnalytical Services Laboratory (BAS Lab) at IMET on an Illumina MiSeq™ (San Diego, CA, USA) using 5 ng DNA from each sample. Primers complementary to the V3-V4 hypervariable region of the bacterial 16s rRNA gene were designed based on those characterized by Klindworth et al.: S-D-Bact-0341-b-S-17 5’- CCTACGGGNGGCWGCAG-3’ and S-D-Bact-0785-a-A-21 5’-GACTACHVGGGTATCTAATCC-3’ (28). Including the Illumina adaptor sequences, the full-length primers were F 5’ TCGTCGGCAGCGTCAGATGTGTATAAGAGACAGCCTACGGGNGGCWGCAG and R 5’ GTCTCGTGGGCTCGGAGATGTGTATAAGAGACAGGACTACHVGGGTATCTAATCC. CLC Genomics Workbench 8 (Version 8.5.1, Qiagen, Germantown, MD, USA) was used to trim, pair, and merge sequences (default parameters used for all functions). They were exported as a merged FASTA file and imported into the Quantitative Insights into Microbial Ecology (QIIME) program (Version 1.9.1) (29) for open reference operational taxonomic unit (OTU) and taxonomic classification using the Silva 128 reference database (30). Rarefaction curves were generated based on observed OTUs. Identify threshold was set to 97%. Representative sequence alignments for each OTU were generated using Python Nearest Alignment Space Termination (PyNAST) (31), and R (Version 3.4.3) (32) was used to generate a bar graph of the bacterial orders, as well as a PCA plot. QIIME was used to generation the rarefaction plot.

#### Statistics

Statistical significance was evaluated using analysis of variance (ANOVA) and Student’s t-test (2-tailed) with a 95% confidence interval.

## Results

### Cobia Trials

Diets 1-4 for both PP (plant protein) and FM (fishmeal) were formulated to contain graded levels of taurine of approximately 0, 0.5%, 1.5% and 5%. PP1-4 contain approximately 3.6%, 3.6%, 3.5%, and 3.2% total wheat gluten, respectively. This calculation includes any wheat gluten present in the wheat flour, estimated to be ∼8% of the total weight. The FM1-4 diets contain an average of 1.8% total wheat gluten. These diets are part of a previously published study performed by our laboratory (33). The EPP3 diet contains approximately 1.2% wheat gluten and 1.5% taurine.

### Growth Data

Growth data for Trial 1 of diets PP1-PP4 are displayed in Figure 1 (A). For this trial, the average initial fish weight was ∼10 g, and fish were maintained on the diets for 8 weeks. Data from Trial 3 with fish of average initial weight ∼18 g fed the EPP3 diet are also included on this graph, as well as averaged growth data for fish fed various fishmeal diets (FM1-FM4). Growth data from Trial 2 (∼120 g initial weight) are shown in Figure 1 (B). The trial continued for 8 weeks and as with the graph for the first trial, data are included for comparison from the EPP3 and FM1-FM4 studies. For both trials, the fishmeal and EPP3 diets trend higher in terms of growth. PP1 appears to be the least favorable diet in terms of growth. It contains the greatest amount of wheat gluten but also contains no taurine.

**Figure 1:**
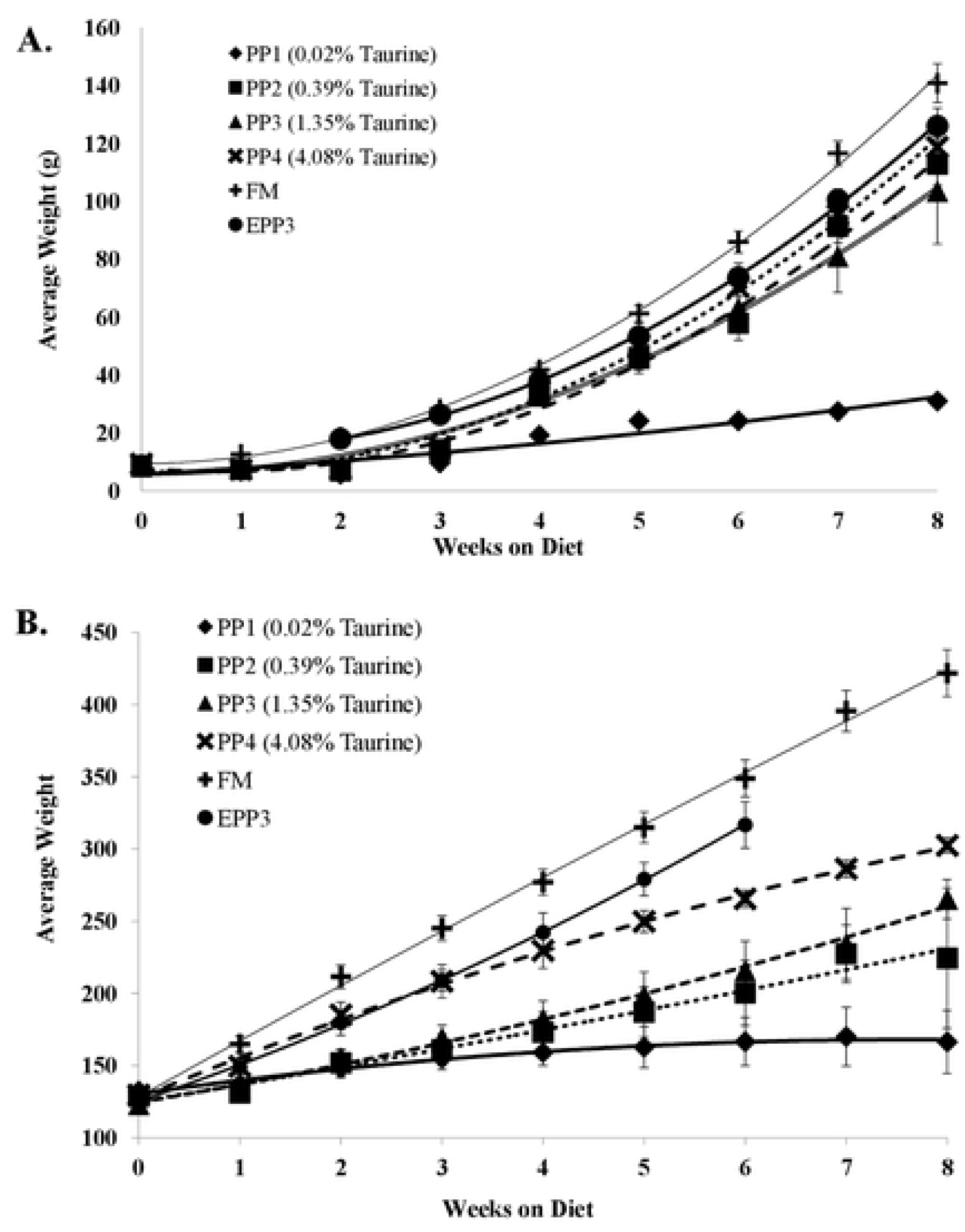
Growth data from 8-week dietary trials in juvenile cobia. Plot A presents data from the first growth trial starting with ∼10 g average weight juveniles for four plant protein based diets (PP1, PP2, PP3, and PP4) with graded levels of supplemental taurine and the average of four fishmeal-based diets (FM, data from Watson et al. 2014) (33). Plot B presents data from the second growth trial starting with ∼120 g average weight juveniles. Included are the growth data for the trial with diet EPP3 starting with ∼18 g average weight juveniles. Plotted are the average tank weights from 3 replicates ± standard deviation.

### Survival and performance characteristics

Performance characteristics for fish fed PP1-PP4 are shown in Table 1. Differences in survival are substantial, with PP4 containing the lowest amount of wheat gluten and highest amount of taurine being the best performer. There were no significant differences among surviving individuals in percent weight gain, feed conversion ratio (FCR), specific growth rate (SGR), total plasma protein concentration, or total bile salt concentration (ANOVA, P>0.05). Due to low survival in several replicates of PP1 and PP2 resulting in 0 individuals in some tanks, these diets were not included in statistical analyses other than survival for the first trial.

**Table 1:** Performance characteristics from Trial 1 with cobia (∼10 g initial weight). Within a row, values that share common superscripts are not significantly different from one another (P>0.05).

| Characteristic | PP1 (0.02% T) | PP2 (0.39% T) | PP3 (1.35% T) | PP4 (4.08% T) |
| --- | --- | --- | --- | --- |
| Survival (%) | 1.96 ± 1.96<sup>a</sup> | 9.80 ± 5.18<sup>a,b</sup> | 9.83 ± 1.97<sup>a,b</sup> | 19.6 ± 7.06<sup>b</sup> |
| Weight Gain (%) | 315.55<sup>3</sup> | 1194.73 ± 21.22<sup>3</sup> | 1155.92 ± 410.70 | 1200.68 ± 214.22 |
| FCR<sup>1</sup> | 1.93<sup>3</sup> | 0.96 ± 0.02<sup>3</sup> | 1.17 ± 0.44 | 0.95 ± 0.11 |
| SGR<sup>2</sup> | 2.54<sup>3</sup> | 4.57 ± 0.03<sup>3</sup> | 4.26 ± 0.72 | 4.53 ± 0.31 |
| Total Plasma Protein (g dL<sup>-1</sup>) | 3.49<sup>3</sup> | 3.40 ± 0.17<sup>3</sup> | 3.35 ± 0.17 | 3.48 ± 0.23 |
| Total Bile Salts (mM) | 40.84<sup>3</sup> | 42.17 ± 0.46<sup>3</sup> | 41.37 ± 1.99 | 41.50 ± 1.36 |

T=Taurine
<sup>1</sup> FCR =feed conversion ratio = (g fed/g gained)
<sup>2</sup> SGR=specific growth rate = $((\ln BW_f - \ln BW_i) * (\text{days of growth trial}^{-1})) * 100$
<sup>3</sup> Not included in statistical analyses due to lack of replicates (survivors)

Performance characteristics from the second trial (∼120 g initial weight) are shown in Table 2. Percent weight gain was significantly lower in diets PP1 and PP2 compared to the other four fishmeal-based diets (p<0.05), but not significantly different from diets PP3 and PP4 (p>0.05). Diet PP1 resulted in significantly lower FCR and SGR than the other diets (p<0.05). Diets PP2, PP3, and PP4 did not result in significantly different SGRs than one another (p>0.05). The compared outcomes from diets PP1 and PP2 containing roughly the same amount of wheat gluten with and without taurine supplementation reinforce the importance of the amino acid for promoting growth and suggest that it may partially alleviate adverse effects resulting from wheat gluten. Total bile salt concentration was not significantly different between any of the 4 dietary treatments (p>0.05).

**Table 2:** Performance characteristics from Trial 2 with cobia (∼120 g initial weight). Within a row, values that share common superscripts are not significantly different from one another (P>0.05).

| Characteristic | PP1 (0.02% T) | PP2 (0.39% T) | PP3 (1.35% T) | PP4 (4.08% T) |
| --- | --- | --- | --- | --- |
| Weight Gain (%) | 23.35 ± 22.82<sup>a</sup> | 80.57 ± 60.24<sup>a</sup> | 130.87 ± 25.20<sup>b</sup> | 133.82 ± 10.83<sup>b</sup> |
| FCR | 6.38 ± 1.49<sup>a</sup> | 2.97 ± 1.17<sup>b</sup> | 1.98 ± 0.19<sup>b</sup> | 2.12 ± 0.23<sup>b</sup> |
| SGR | 0.57 ± 0.12<sup>a</sup> | 1.31 ± 0.26<sup>b</sup> | 1.47 ± 0.19<sup>b</sup> | 1.51 ± 0.09<sup>b</sup> |
| Total Bile Salts (mM) | 38.76 ± 7.40 | 28.37 ± 1.94 | 36.30 ± 5.69 | 28.75 ± 3.22 |

T=Taurine
<sup>1</sup> FCR =feed conversion ratio = (g fed/g gained)
<sup>2</sup> SGR=specific growth rate = $((\ln BW_f - \ln BW_i) * (\text{days of growth trial}^{-1})) * 100$

For the EPP3 diet in the third trial (initial weight ∼18 g), weight gain, FCR, and SGR were much improved from the first two trials, and were an improvement over all previous plant protein diets tested in our laboratory. Performance characteristics included weight gain of 1673.52% ± 192.75, FCR of 1.22 ± 0.04, and SGR of 3.42 ± 0.13. The EPP3 diet contains the lowest gluten content of all diets tested in this study (1.2%).

### Wheat gluten: Survival and pathology

Figure 2 presents survival curves based on percent wheat gluten for fish fed diets PP1-PP4 (Trial 1), EPP3 (Trial 3), or fishmeal-based diets FM1-FM4 (averaged) from a previous study (33). These data suggest that wheat gluten inclusion may be deleterious to cobia at this early stage of development, though it may be tolerated at later stages. Histological analysis of the proximal intestinal epithelium showed no distinct differences for fish fed the PP1-PP4 and FM1-FM4 diets. However, pathological effects in the distal intestinal region cannot be ruled out. No histological analysis was performed on the EPP3 fish. Survival was approximately 100% for fish in Trial 2 fed the PP1-PP4 diets and in Trial 3 with the EPP3 diet. Cannibalism was not observed to be a contributing factor to the low survival in Trial 1, and dead individuals were promptly removed from the tanks so as to not be a nutritional/taurine source for remaining fish.

**Figure 2:**
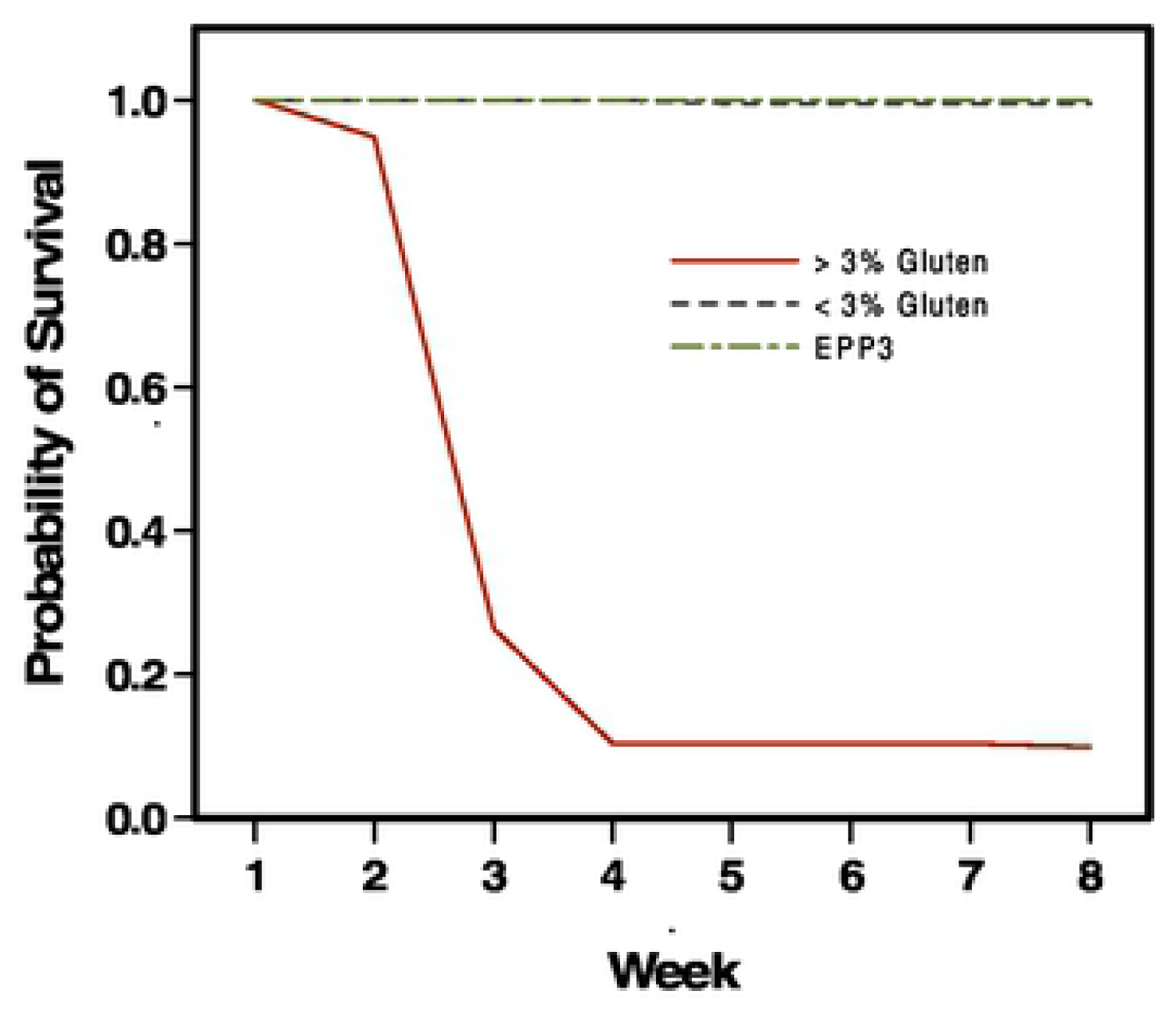
Dietary wheat gluten level affects survival of young cobia. For Trial 1 fish (growth data shown in Figure 1(A)), probability of survival is graphed as a function of dietary gluten. For fish fed PP1-PP4, (> 3% gluten) or FM1-FM4 (< 3% gluten) (33), survival data from the diet subtypes (1,2,3, and 4) are averaged. Also shown is survival data from fish fed EPP3 (1.2% gluten).

### Muscle and liver characteristics

Fillet and liver characteristics from the second and third trials are shown in S6 Table: Fillet and liver characteristics from the second and third cobia growth trials. Fillet water content was highest in fish fed the PP1 diet with a gradual reduction in fillet water content as dietary taurine level increased. Hepatosomatic index show showed a similar trend of reduction as dietary taurine level increased. Fillet yield, liver water, fillet taurine, and liver taurine contents all showed increasing trends with increasing dietary taurine levels.

### Plasma analysis

Results of the plasma analysis are shown in Table 3. Plasma water content significantly decreased as dietary taurine level increased (p<0.05). Concentrations of glucose, calcium, phosphorus, and magnesium ions, as well as overall osmolality, are indicators of health status. Total protein levels also often correlate with nutritional status and general health (34), (35). Triglycerides are an energy substrate, and a fasting study in European sea bass demonstrated their potential as a marker for nutritional condition (36). AST (aspartate aminotransferase) and ALP (alkaline phosphatase) are indicators of liver health and function (37).

**Table 3:** Plasma measures from fish from the second and third cobia growth trials. Values represent the mean ± standard error for three fish per dietary treatment. Within a row, values that share common superscripts are not significantly different from one another (p >0.05).

| Plasma Component | PP1<br>(0.02% T)<sup>1</sup> | PP2<br>(0.39% T)<sup>1</sup> | PP3<br>(1.35% T)<sup>1</sup> | PP4<br>(4.08% T)<sup>1</sup> | EPP3<br>(1.05% T)<sup>2</sup> |
| --- | --- | --- | --- | --- | --- |
| Water Content (%) | 96.75 ±<br>0.19<sup>a</sup> | 94.75 ±<br>0.21<sup>b</sup> | 95.19 ±<br>0.29<sup>a,b</sup> | 94.73 ±<br>0.13<sup>b</sup> | 94.94 ±<br>1.21<sup>a,b</sup> |
| Osmolality (Osm L<sup>-1</sup>) | 329.17 ±<br>5.57 | 343.00 ±<br>5.57 | 325.17 ±<br>18.94 | 336.33 ±<br>4.60 | 332.84 ±<br>9.65 |
| Taurine (nmol ml<sup>-1</sup>) | 421.25 ±<br>41.03<sup>a</sup> | 565.79 ±<br>17.91<sup>a</sup> | 658.47 ±<br>50.92<sup>b</sup> | 723.86 ±<br>89.62<sup>b</sup> | 601.29 ±<br>47.97<sup>a,b</sup> |
| Albumin (g dL<sup>-1</sup>) | 0.53 ±<br>0.12<sup>a</sup> | 0.70 ±<br>0.06<sup>a,b</sup> | 0.80 ±<br>0.00<sup>a,b</sup> | 0.87 ±<br>0.03<sup>b</sup> | 0.52 ±<br>0.07<sup>a</sup> |
| Total Bilirubin (mg dL<sup>-1</sup>) | 0.40 ±<br>0.10<sup>a</sup> | 0.30 ±<br>0.00<sup>a</sup> | 0.23 ±<br>0.03<sup>a</sup> | 0.23 ±<br>0.03<sup>a</sup> | 0.12 ±<br>0.00<sup>b</sup> |
| Calcium (mg dL<sup>-1</sup>) | 9.70 ± 0.88 | 10.90 ±<br>0.61 | 11.10 ±<br>0.10 | 11.37 ±<br>0.27 | 11.69 ±<br>0.30 |
| Cholesterol (mg dL<sup>-1</sup>) | 49.33 ±<br>12.17<sup>a</sup> | 68.33 ±<br>1.20<sup>a,b</sup> | 80.33 ±<br>3.18<sup>b</sup> | 79.33 ±<br>4.26<sup>a,b</sup> | 46.59 ±<br>11.43<sup>a</sup> |
| Creatine Kinase (U L<sup>-1</sup>) | 96.67 ±<br>46.77 | 386.67 ±<br>276.01 | 301.33 ±<br>81.63 | 552.00 ±<br>113.15 | nd |
| Creatinine (mg dL<sup>-1</sup>) | 0.17 ± 0.03 | 0.20 ± 0.00 | 0.17 ± 0.03 | 0.17 ±<br>0.03 | nd |
| Glucose (mg dL<sup>-1</sup>) | 48.00 ±<br>4.36<sup>a</sup> | 51.33 ±<br>3.48<sup>a</sup> | 48.33 ±<br>3.92<sup>a</sup> | 41.67 ±<br>3.93<sup>a</sup> | 109.13 ±<br>19.13<sup>b</sup> |
| Phosphorous (mg dL<sup>-1</sup>) | 7.13 ±<br>1.34<sup>a</sup> | 8.37 ±<br>0.69<sup>a</sup> | 8.87 ±<br>0.38<sup>a</sup> | 9.33 ±<br>0.23<sup>a</sup> | 13.82 ±<br>1.78<sup>b</sup> |
| Magnesium (mg dL<sup>-1</sup>) | 2.37 ± 0.22 | 1.97 ± 0.07 | 2.13 ± 0.03 | 2.23 ±<br>0.08 | 2.77 ±<br>0.18 |
| Triglycerides (mg dL<sup>-1</sup>) | 66.67 ±<br>27.43 | 112.00 ±<br>19.67 | 91.00 ±<br>36.12 | 109.67 ±<br>20.96 | 75.73 ±<br>17.35 |
| Sodium (mmol L<sup>-1</sup>) | 169.10 ±<br>3.56 | 178.67 ±<br>2.89 | 177.33 ±<br>2.32 | 174.83 ±<br>3.89 | nd |
| Potassium (mmol L<sup>-1</sup>) | 8.78 ± 1.09 | 8.07 ± 0.36 | 8.92 ± 0.53 | 9.35 ±<br>0.23 | nd |
| Chloride (mmol L<sup>-1</sup>) | 169.97 ±<br>1.93 | 173.97 ±<br>2.08 | 168.83 ±<br>0.88 | 169.43 ±<br>3.85 | nd |

T=Taurine
<sup>1</sup> UCLA DLAM analysis
<sup>2</sup> NOAA-NWFSC analysis by A.W. (excluding water content, osmolality, and taurine)

Plasma taurine levels were significantly lower in diets PP1 and PP2 than the other diets (p<0.05). Plasma cholesterol, phosphorous, and albumin all showed similar trends of significantly increasing with dietary taurine level (p<0.05). Lower creatine kinase levels can be indicative of muscle wasting (38). Though biochemical markers of liver failure are not well characterized in fish, elevated bilirubin is a classic sign (39). Stress on the liver may be a result of insufficient taurine required for bile acid conjugation or a manifestation of gluten-induced enteritis (40). The substantial difference in plasma glucose levels between PP1-PP4 and EPP3 is likely an artifact of MS-222 and not a barometer of health status (41).

Figure 3 shows H&E-stained liver sections from Trial 2 fish fed PP1 (Panel A) or PP4 (Panel B). Supplementation with taurine prevents pervasive steatosis. This effect was also seen in fish fed FM1 vs. FM4 (data not shown). Taurine supplementation may have therapeutic potential in the treatment of nonalcoholic fatty liver disease, suggesting that supplementation may be of value even in cases where there is not an underlying deficiency (42).

**Figure 3:**
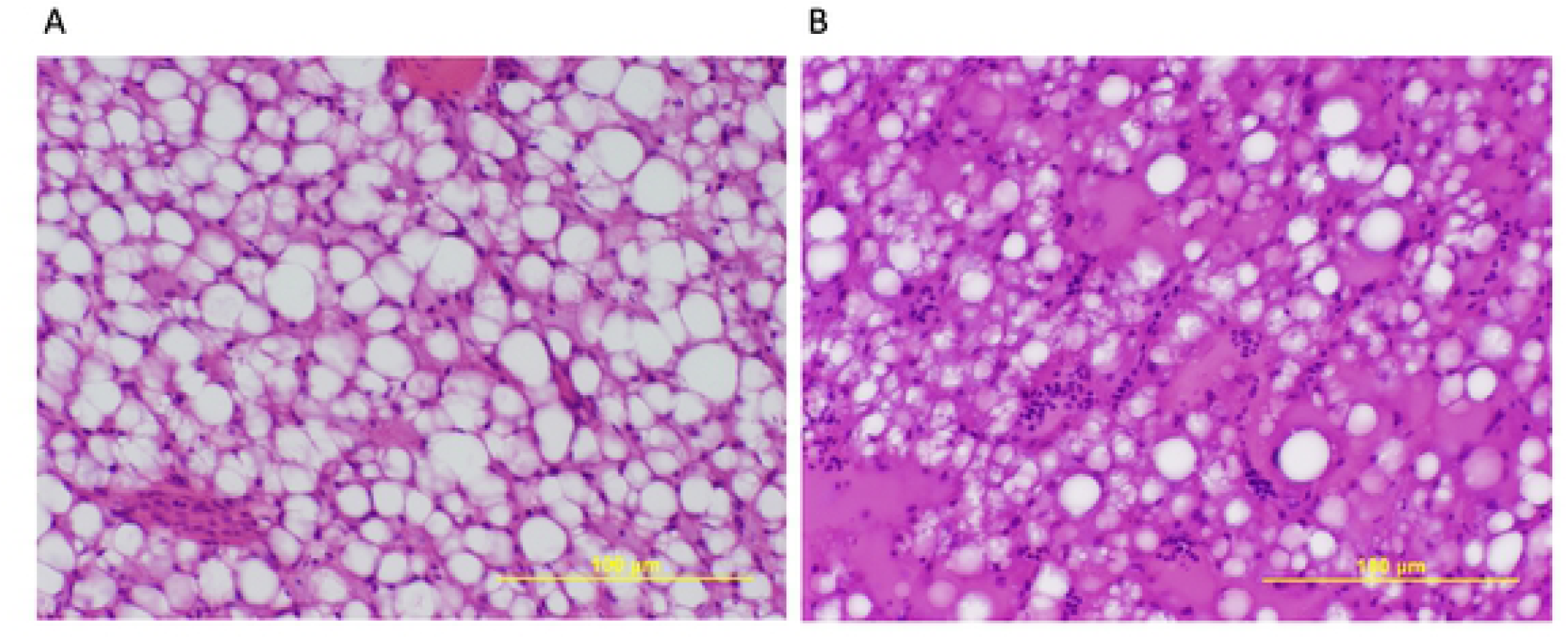
Taurine supplementation mediates hepatic steatosis. Histological sections from fish were stained with H&E and microscopically examined. Panel A presents a representative section from fish fed PP1 (0.02% taurine). Panel B presents a representative section from fish fed PP4 (4.08% taurine). Pervasive lipidosis is apparent in Panel A.

### Detection of immune factors capable of binding to wheat gluten

Suspecting that wheat gluten might be a contributing factor to poor growth performance, we assayed in Trial 2 fish plasma an immune response to gliadin, a potentially immunogenic component of wheat gluten. Standard methods for detecting specific antibody responses to wheat gluten in human celiac patients cannot be utilized as they are designed to detect IgA. Cobia and other teleosts have only three classes of immunoglobulins: IgM, IgT, and IgD, and there are currently no reagents to detect these immunoglobulins in cobia (43). It is not known if antibodies designed to detect these adaptive factors have cross-reactivity to these immune components in cobia.

To detect plasma factors capable of binding to gliadin, varying concentrations of the protein as well as a fixed amount of eIF4E-1A from the dinoflagellate *A. carterae* were subjected to immunoblotting. eIF4E-1A, a translation protein, was used as a control for loading, transfer efficiency, and specificity of binding. After electroblotting, membranes were pre-incubated in blocking buffer only or blocking buffer containing plasma. The 30-40 kDa bands visible on the immunoblots correspond to a range of α/β and γ gliadins (44). These gliadin bands are less visible over the course of 2-fold dilutions (Figures 4 and 5).

**Figure 4:**
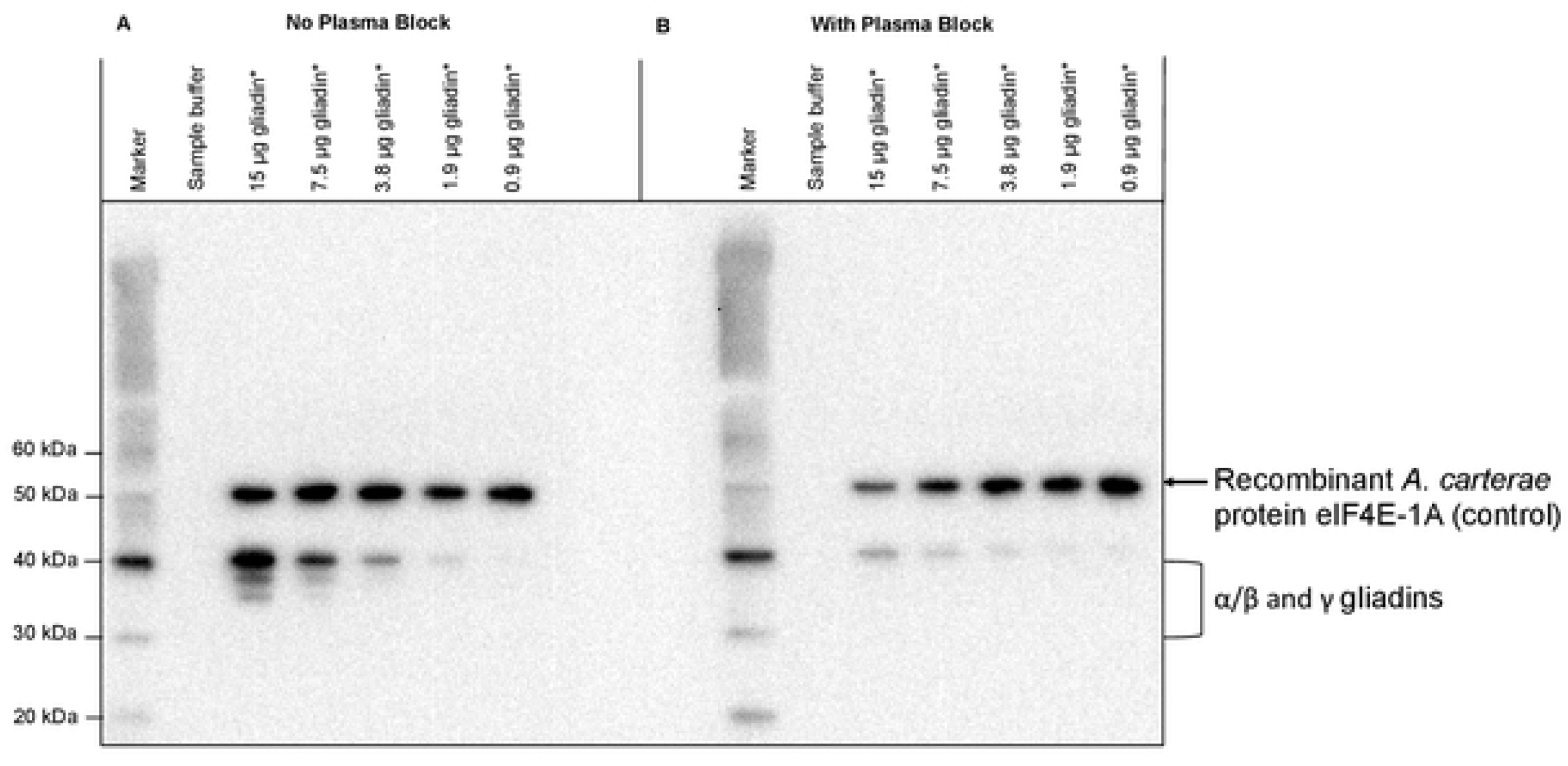
Factor(s) that specifically bind gliadin are present in plasma of fish fed plant-based diets containing 3.2-3.6% wheat gluten. Immunoblotting of gliadin without pre-incubation of plasma (A) or with pre-incubation with plasma from fish fed diets containing 3.2-3.6% wheat gluten from Trial 2 (B) demonstrates the ability of factor(s) in plasma to bind to gliadin (30-40 kDa) and inhibit binding of anti-gliadin polyclonal antibody. Each lane also contains 431 ng recombinant A. *carterae* protein eIF4E-1A (50 kDa).

**Figure 5:**
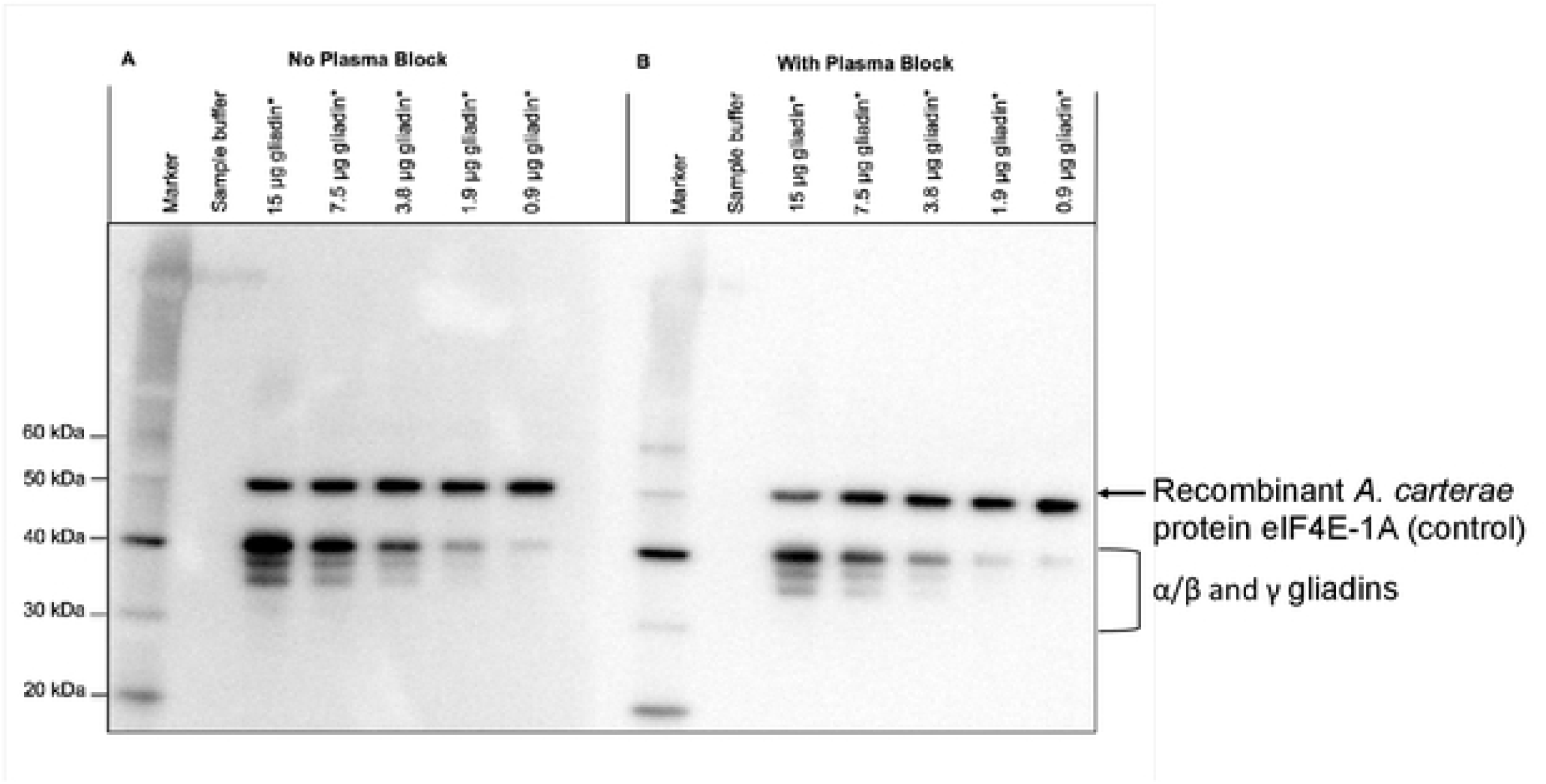
Factor(s) that bind gliadin are absent in plasma from wild cobia with no dietary gluten exposure. Immunoblotting of gliadin without pre-incubation of plasma (A) or with pre-incubation with plasma (B). No factors capable of binding to gliadin (30-40 kDa) and inhibiting binding of anti-gliadin polyclonal antibody are detectable. Each lane also contains 431 ng recombinant *A. carterae* protein eIF4E-1A (50 kDa).

Pre-incubation with plasma from cobia fed diets containing 3.2-3.6% wheat gluten diminished binding of an anti-gliadin polyclonal antibody, even at the highest concentration of gliadin (Figure 4). A comparison of slopes from linear regressions based on densitometry data for “No Plasma Block” vs. “With Plasma Block” relatively quantifies this difference in signal as approximately 10-fold. A similar dot blot experiment with gliadin but no eIF4E-1A confirmed the results of the western blot (not shown). Similar results were obtained using plasma from fish fed diets containing 1.5-1.9% wheat gluten, suggesting that a small amount is sufficient to mobilize a response (not shown).

Pre-incubation with plasma from wild-caught cobia with no dietary exposure to wheat gluten does not diminish binding of an anti-gliadin polyclonal antibody, even at the lowest concentrations of gliadin (Figure 5). For all experiments, there was no reduction in eIF4E-1A signal as a result of pre-incubation with plasma, demonstrating the specificity of plasma factors for gliadin. There was also no detection of a component capable of binding gliadin found in the plasma of fish fed the FM1-4 diets (data not shown).

### European sea bass

#### Growth data

Growth rates were equivalent (p>0.05) for the fish fed plant-based diets with or without added wheat gluten (4%). The study commenced with fish of an average weight of ∼25 g and continued for 6 months. The growth curve is shown in Figure 6.

**Figure 6:**
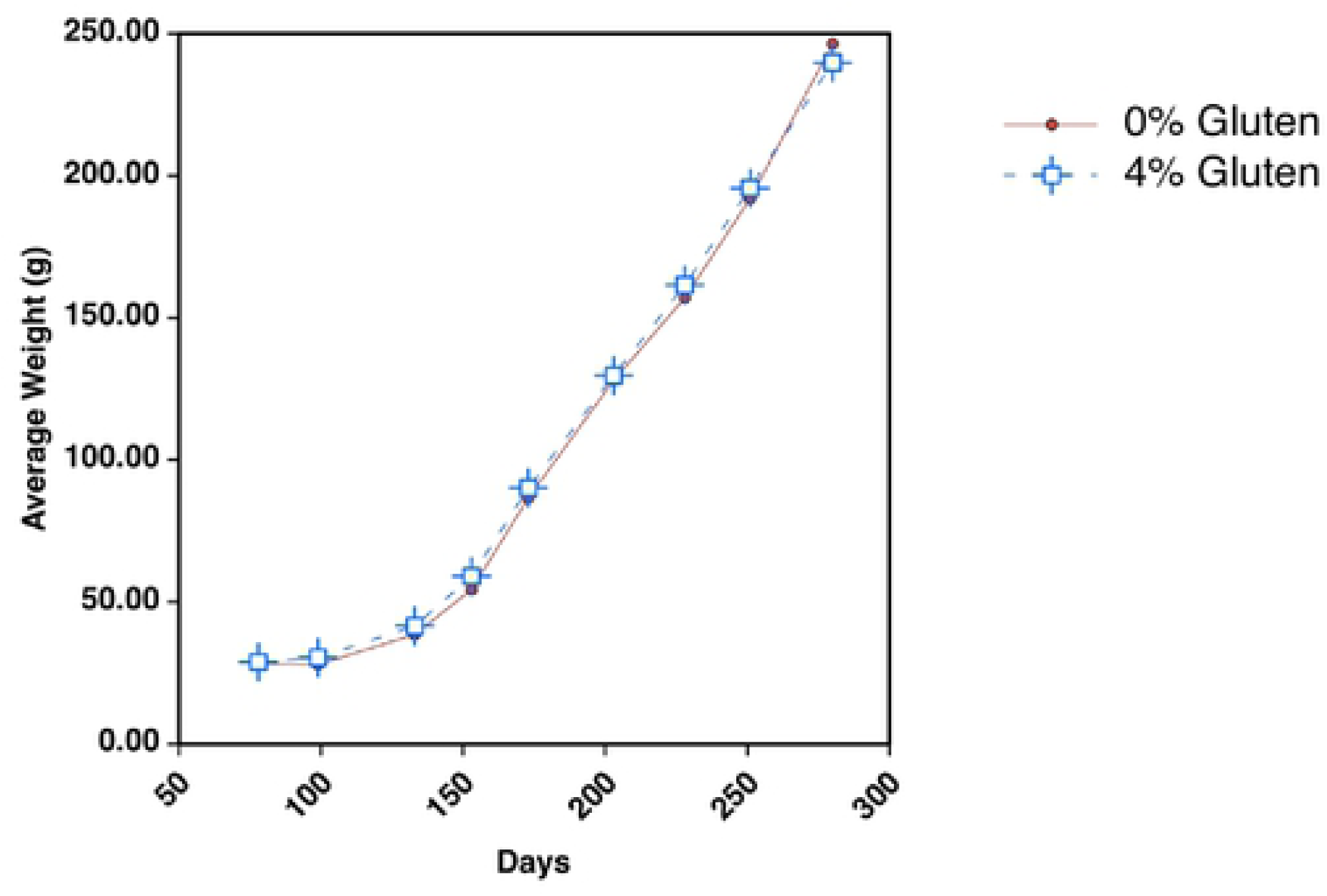
Dietary inclusion of 4% wheat gluten does not affect growth of European sea bass. European sea bass were fed plant-based diets containing 0 or 4% wheat gluten starting at an average weight of ∼25 g. The study continued for 6 months, and growth rates were similar throughout the study (p>0.05).

#### Plasma analysis

As shown in Table 4, analysis of common plasma analytes using a bioanalyzer revealed no differences in levels between the 0 or 4% wheat gluten dietary groups with the exception of calcium and AST (aspartate aminotransferase) (p<0.05). Plasma calcium is higher in the group fed 4% wheat gluten, whereas plasma AST levels are lower.

**Table 4:** Levels of common plasma parameters for European sea bass fed diets with 0 or 4% wheat gluten are similar in value except for levels of calcium and AST.

| Plasma Component | 0 Wheat Gluten | 4% Wheat Gluten |
| --- | --- | --- |
| Glucose (mg/dL) | 120.727 ± 27.335 | 113.874 ± 32.011 |
| **Calcium (mg/dL)** | **\*11.045 ± 0.465** | **\*11.75 ± 0.567** |
| Magnesium (mg/dL) | 3.545 ± 0.398 | 3.7 ± 0.25 |
| Phosphorus (mg/dL) | 7.909 ± 0.76 | 8.125 ± 0.784 |
| Triglycerides (mg/dL) | 410.364 ± 77.747 | 388 ± 129.836 |
| Cholesterol (mg/dL) | 199.545 ± 33.074 | 206.75 ± 43.702 |
| ALP (U/L) | 24.091 ± 1.486 | 25.125 ± 1.511 |
| **AST (U/L)** | **\*47.8 ± 28.558** | **\*35.25 ± 17.484** |
| Osmolality (mOsmol/kg) | 363 ± 5.773 | 368.917 ± 6.851 |
| Total Protein (g/dL) | 4.291 ± 0.241 | 4 ± 0.525 |

\*Statistically different between diets
ALP=Alkaline phosphatase
AST=Aspartate aminotransferase

#### Body and tissue weights

Whole body and tissue weights are similar between fish fed either 0 or 4% wheat gluten with the exception of the mid-intestine (p<0.05). This is not a typical place for major changes to the intestine to manifest, so this is an interesting result. These data are summarized in S7 Table: Body and tissue weights for fish fed 0 or 4% wheat gluten are similar with differences manifesting in the mid-intestine. Hepatosomatic indices were an average of 0.01 for both dietary groups.

#### Plasma taurine analysis

Though there were minimal differences between the diets for levels of most common plasma analytes, a separate analysis of plasma taurine levels by HPLC showed that levels were approximately twice as high in the fish fed the diet containing wheat gluten (Figure 7). One possible explanation is that the fish are producing or sequestering more taurine to counter some effect induced by the wheat gluten.

**Figure 7:**
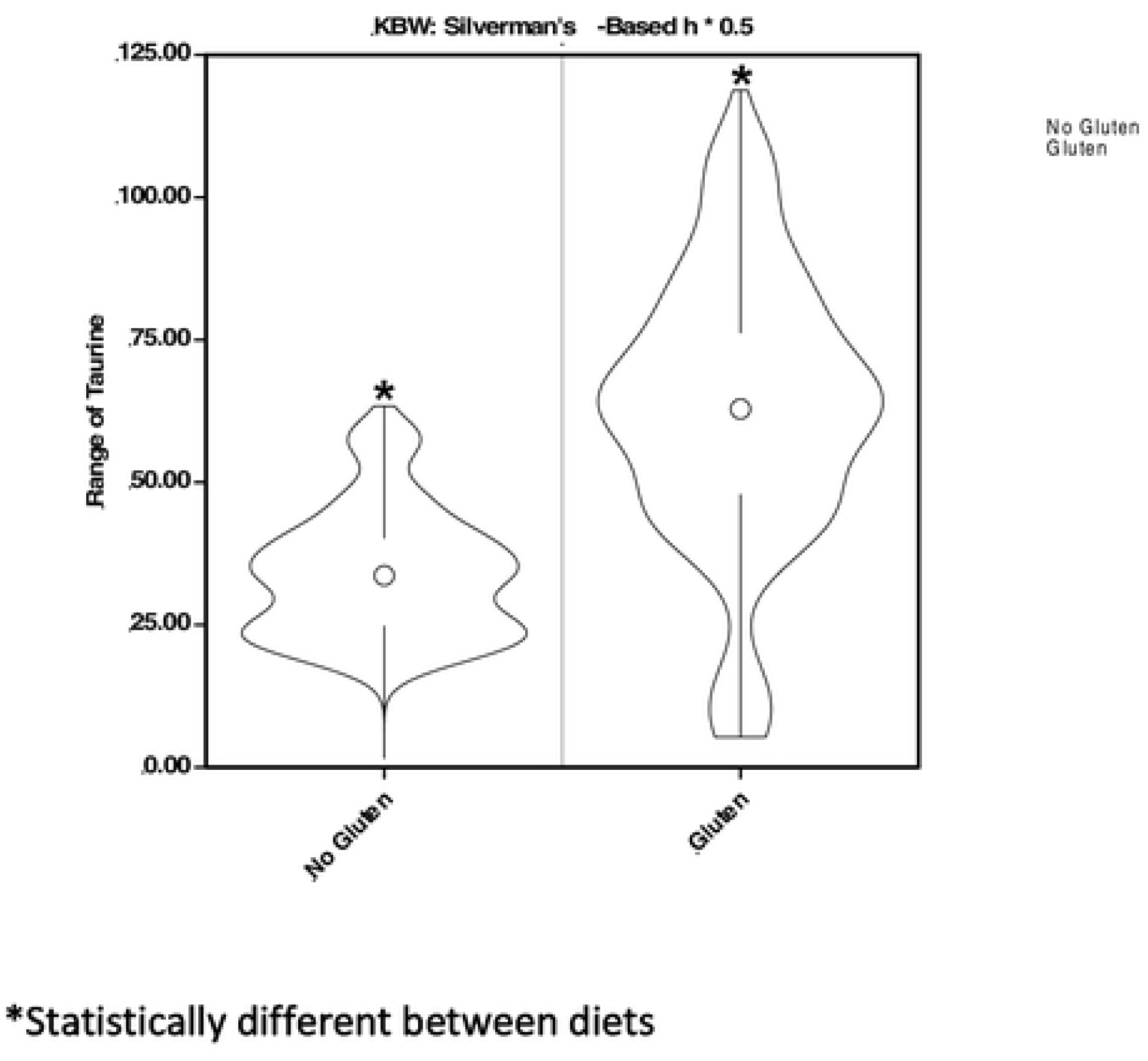
4% dietary wheat gluten substantially raises plasma taurine levels in European sea bass. Taurine levels in plasma were measured using HPLC. Average concentrations of plasma taurine for fish fed the 0 or 4% wheat gluten diets are 33.82 ± 7.133 or 62.515 ± 15.719 nmol/mL, respectively.

#### Gliadin immunoblotting

To detect plasma factors capable of binding to gliadin, varying concentrations of the protein as well as a fixed amount of eIF4E-1A from the dinoflagellate *A. carterae* were subjected to immunoblotting. eIF4E-1A, a translation protein, was used as a control for loading, transfer efficiency, and specificity of binding. After immunoblotting, membranes were pre-incubated in blocking buffer only or blocking buffer containing plasma. In the immunoblot, bands are visible in the 30-40 kDa range corresponding to various α/β and γ gliadins (Figure 8) (44). These gliadin bands are less visible over the course of 2-fold dilutions. There does not appear to be a component in European sea bass plasma produced in response to dietary wheat gluten that is capable of binding gliadin. This would be evidenced by decreased binding of the primary antibody, and subsequently the secondary signal antibody. This is in contrast to data presented in Chapter 2 showing that juvenile cobia produce a plasma factor capable of binding gliadin when they are fed a diet containing 3.2-3.6% wheat gluten.

**Figure 8:**
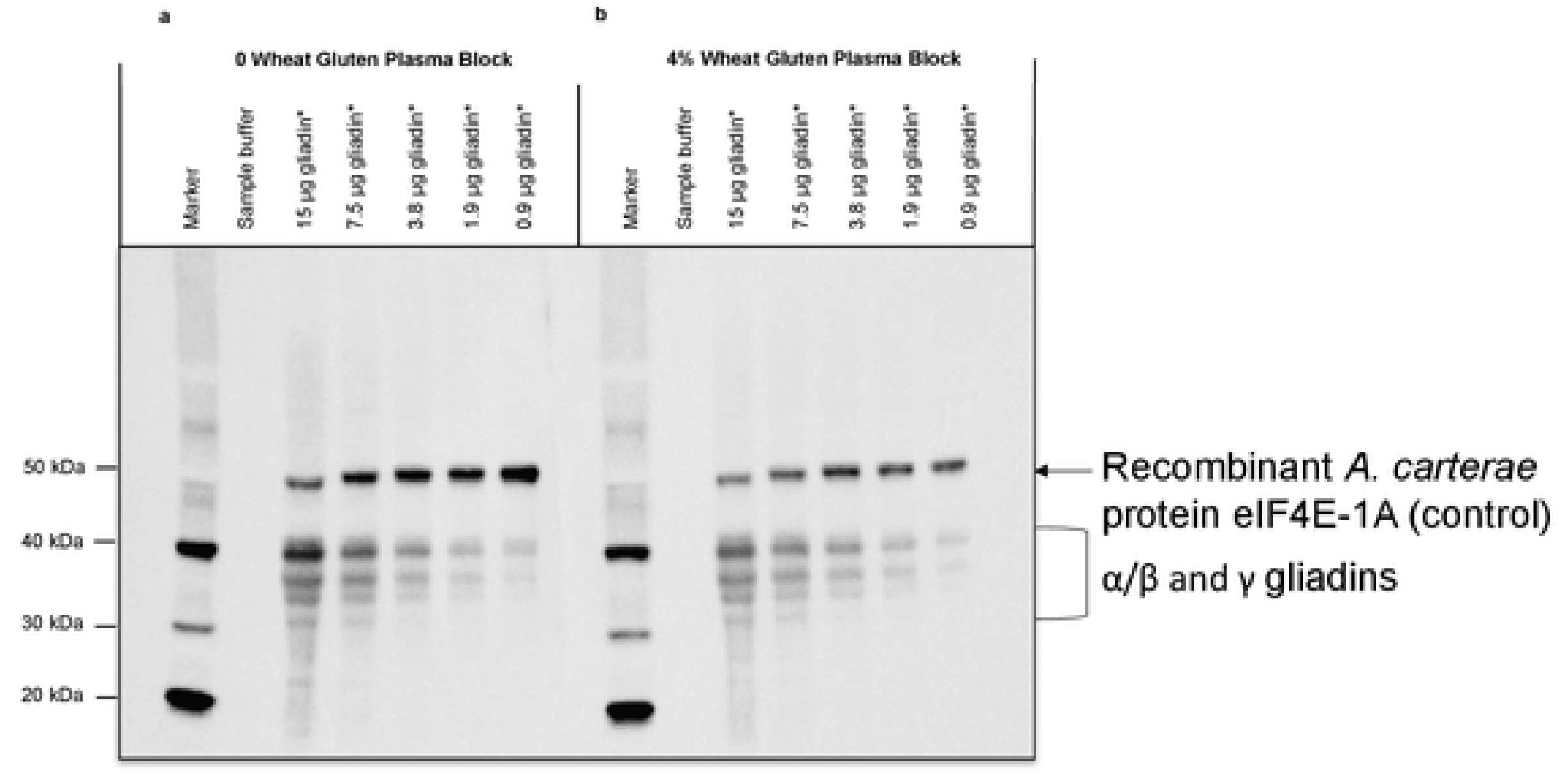
Plasma factors capable of binding gliadin are not detectable in European sea bass fed 4% wheat gluten. Western blot analysis of gliadin without pre-incubation of plasma (a) or with pre-incubation with plasma (b). No factors capable of binding to gliadin (30-40 kDa) and inhibiting binding of anti-gliadin polyclonal antibody are detectable. Each lane also contains 431 ng recombinant *A. carterae* protein eIF4E-1A (50 kDa).

#### Ig and IgM

We used immunoblotting to assay for IgT and IgM to see if dietary inclusion of wheat gluten alters their levels in plasma. The results shown in Figure 9 suggest that levels do not change in response to dietary wheat gluten. Any immune response mounted against gliadin or some other component of wheat gluten is not likely due to an adaptive response, in this case. The size of ∼73 kDa for the heavy chains of the antibodies is similar to the size of ∼78 kDa detected by Picchietti et al. (45).

**Figure 9:**
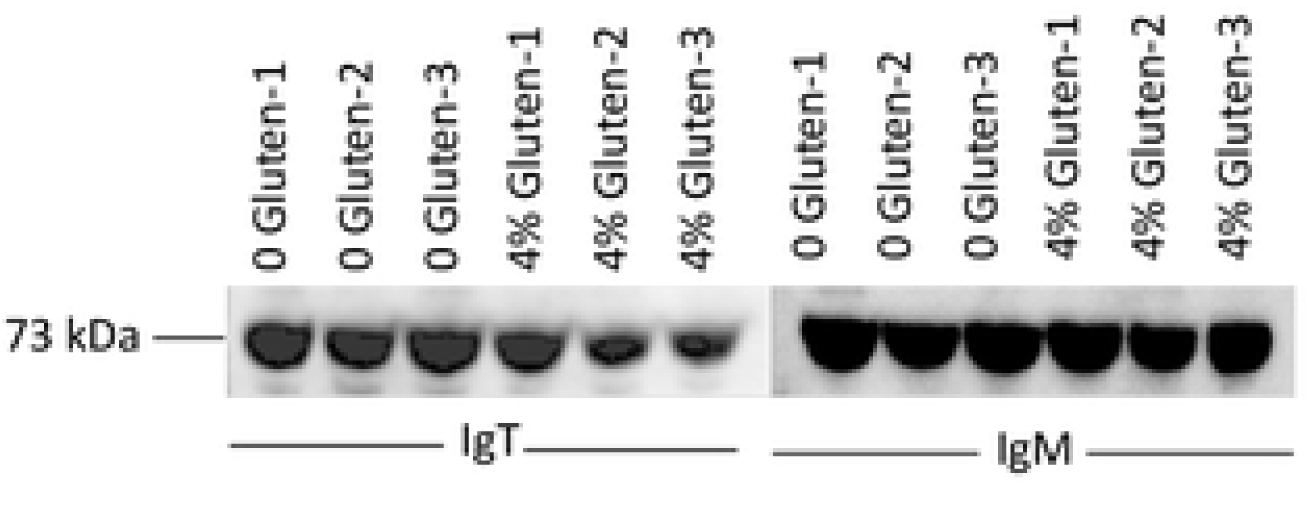
Dietary wheat gluten does not induce changes to levels of plasma IgT or IgM. IgT and IgM were measured using immunoblotting, and their levels do not appear to be altered by 4% dietary wheat gluten.

#### Intestinal microbiome analysis

To characterize the microbiome of the water, feed, and intestinal sections of the European sea bass, 16S rRNA gene analysis was performed using the MiSeq platform. Figure 10 shows a bar graph of order-level taxonomic abundance. Proteobacteria dominate in the tank water. The “cyanobacteria” in feed are most likely chloroplasts from plant ingredients. This is also true for “cyanobacteria” in the pyloric caeca, which likely corresponds to undigested feed despite the fact that food was withheld from these fish for 24 hours before tissue sampling. For fish consuming no dietary gluten (Tank 6-11), the predominant phylum for all intestinal sections analyzed (pyloric caeca, anterior intestine, mid-intestine, and posterior intestine) is Proteobacteria. The same is true for the intestinal sections of the fish fed 4% gluten (Tank 6-12), but there is a greater diversity of predominant orders of Proteobacteria, and Bacteroidetes presents in the mid- and posterior intestines.

**Figure 10:**
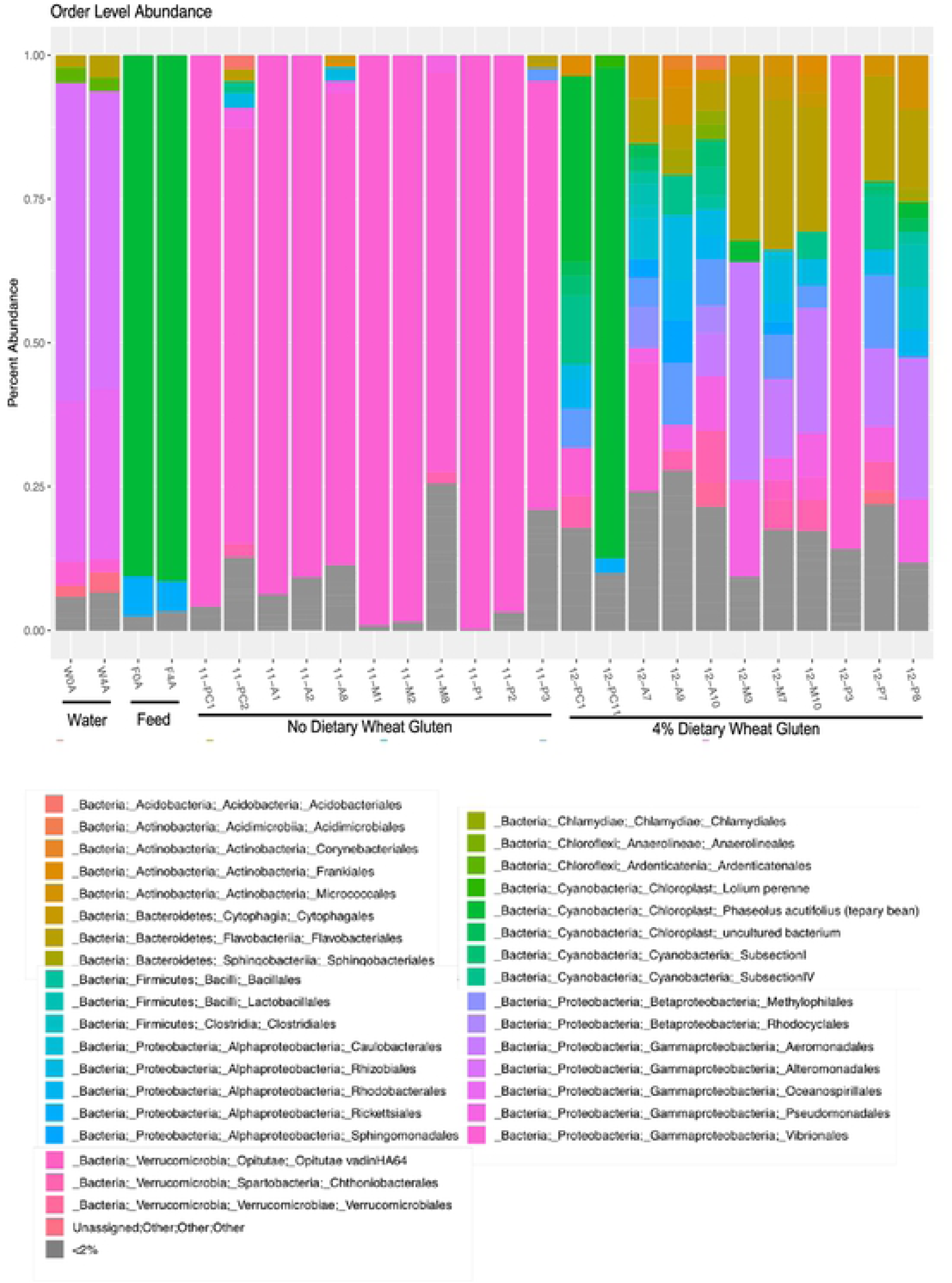
European sea bass fed 4% wheat gluten as part of a plant-based diet exhibit a greater diversity of predominant taxonomic orders across the intestine. DNA extracted from water, feed, and various sections of the intestine for the two diet groups underwent *16S* rRNA gene analysis using the MiSeq platform to characterize the microbial landscape. The addition of 4% wheat gluten to a plant-based diet dramatically shifts the intestinal microbiome of European sea bass.

Figure 11 shows an alpha-diversity rarefaction curve based on OTU data. The PCA (principal component analysis) plot in Figure 12 shows that samples cluster by absence or presence of dietary wheat gluten. The ellipse shows the 95% confidence interval.

**Figure 11:**
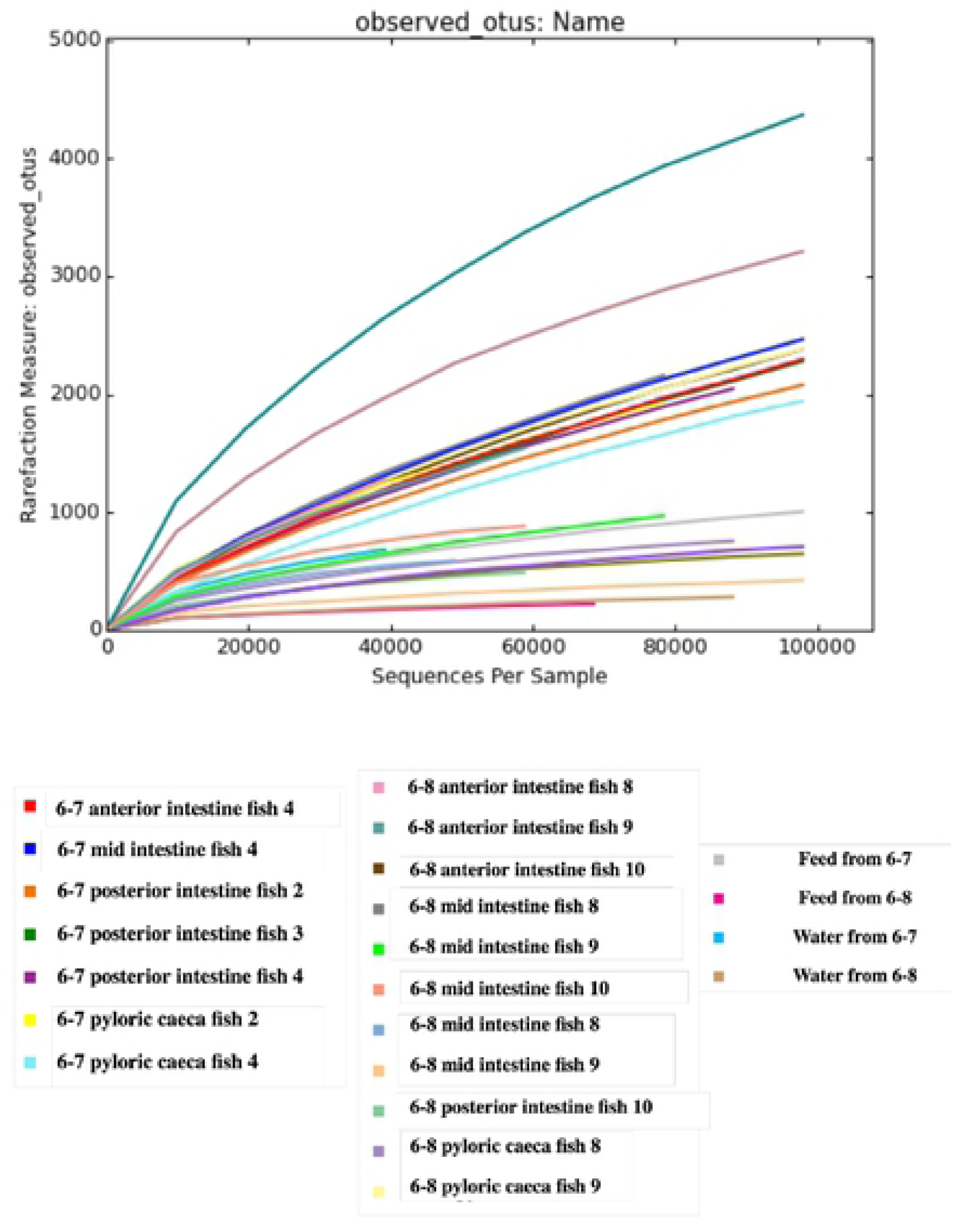
Species richness for each microbiome sample. Species richness of the microbiome samples is shown in a rarefaction curve of observed OTUs vs. sequences per sample.

**Figure 12:**
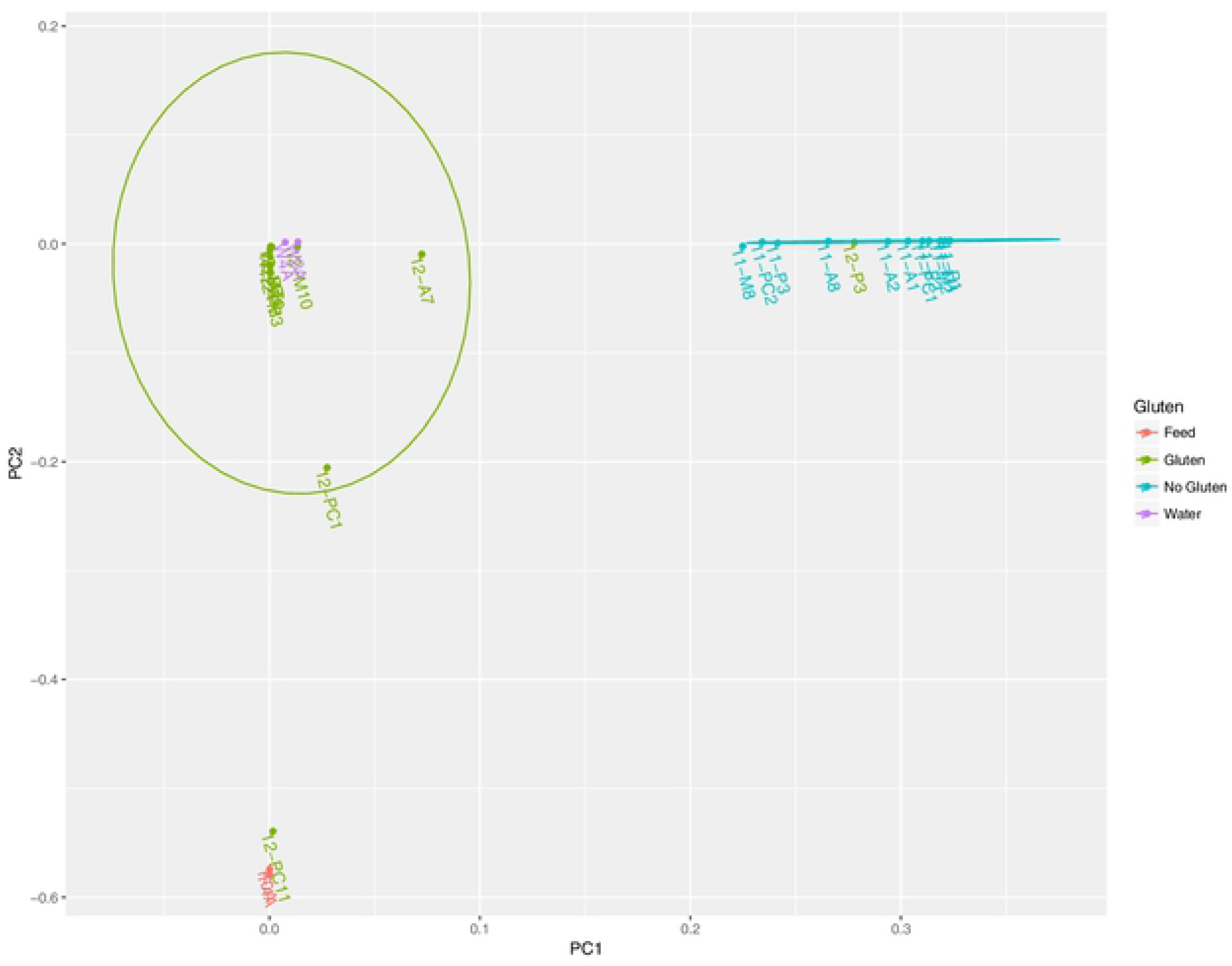
Samples cluster by absence or presence of dietary wheat gluten in a PCA plot. Samples cluster by absence or presence of dietary wheat gluten. The ellipse shows the 95% confidence interval.

## Discussion

Overall, in the cobia study, dietary wheat gluten negatively impacted growth and survival in juvenile cobia, and taurine supplementation appeared to partially ameliorate those effects. Negative effects observed in the first trial are not likely due to a poor cohort as other fish from this brood used for separate feeding trials grew well on both fishmeal-based formulations and commercial feeds in the IMET ARC facility. Palatability issues with plant protein diets have been well described for many species, and it is known that taurine has the potential to serve as a feed attractant due to its small nitrogenous structure (46, 47). Increased feed palatability rather than solely physiological benefits from taurine supplementation may have contributed to the differences in growth and survival during the first trial. Concerns over palatability and the overall formulation prompted the second trial, which was initiated with a size that readily accepted and performed well on plant protein formulations. Although growth and feed conversion in the second trial were lower than for some of the other plant protein formulations such as EPP3, survival was 100% in all treatments, a significant improvement over the first trial. As with the first trial, there was a significant increase in performance with increasing dietary taurine, indicating that taurine is promoting growth and is possibly remediating the negative impacts of this formulation.

The fact that levels of taurine and wheat flour are similar between the high-performing EPP3 diet and the poorer performing PP3 diet suggests that 2.2% added wheat gluten is contributing to a reduction in performance characteristics. There is the possibility that the soy protein concentrate HP300 in EPP3 is greatly enhancing growth characteristics, but trials with another diet, EPP2, point to the gluten as a critical component. EPP2 has an identical formulation to PP3, except EPP3 contains no added wheat gluten with the exception of what is introduced from the wheat flour itself (which is in slightly greater proportion). EPP2 contains roughly 1.8% wheat gluten, and as was the case with EPP3, fish consuming this diet outperformed fish fed the other plant protein-based diets.

The identity of the plasma factor(s) capable of binding gliadin cannot be determined without the necessary reagents to isolate non-specific and specific immune factors in cobia. It is probable that constituents of the cobia innate immune system recognize gliadin, the potentially pathogenic component of wheat. Enzyme-solubilized wheat gluten extracts have been shown to initiate the alternative complement pathway, and it has been suggested that similar to some pathogens, gliadin may be capable of binding to toll-like receptors and initiating an innate immune response (48, 49).

Wheat gluten has been shown to have negative impacts on humans, often but not always affiliated with celiac disease, and other mammalian models (50, 51), (52, 53). It is possible that IgT, the primary mucosal antibody in teleosts, might play a similar role to human IgA. In gluten-sensitive humans, IgA mediates the transport of gliadin across the gastrointestinal epithelium and induces an adaptive immune response (54, 55). Without purified antibodies against cobia IgM or IgT, the role of adaptive immunity cannot be assessed. However, it is clear that ingestion of wheat gluten triggers the mobilization or production of plasma factors capable of binding gliadin, be they innate or adaptive.

The glutamine to glutamate deamidation reaction in gliadin catalyzed by tTG (tissue transglutaminase) is instrumental in gluten-induced pathology. During transamidation, another reaction catalyzed by tTG, the reaction of glutamine with lysine or lysine methyl ester abrogates the T-cell mediated immunotoxic effects of gliadin in the intestine (56). It is possible that the success of wheat gluten incorporation into many fish feeds is due in part to the addition of lysine favoring the transamidation rather than deamidation of gluten catalyzed by tTG. Zebrafish fed a diet containing 60% wheat gluten without adequate lysine supplementation had compromised growth (57). A low pH environment favors deamidation over transamidation, which offers a possible explanation for why gluten may be less tolerated in the gastrointestinal tract of carnivorous fish compared to omnivorous fish (58, 59).

Taurine is present in several tissues and possesses powerful anti-inflammatory properties. It mitigates inflammation by scavenging free radicals, ameliorating oxidative injury, and modulating the immune response (60). Intestinal enteritis has been observed in several species with specific plant protein inclusion such as soy ingredients in salmon and carp (61, 62). Wheat gluten may have triggered inflammation in the cobia intestinal tract, leading to the proliferation of circulating plasma factor(s) capable of binding gliadin.

Previous studies have demonstrated the ability of taurine to mitigate degeneration of the intestinal mucosa and inflammatory bowel disease. It is possible that taurine supplementation of the cobia diet may partially counteract intestinal inflammation caused by dietary ingredients such as wheat gluten (60, 63). Taurine may also offer some protection in the gut against deamidated gluten (64). Unfortunately, samples of the distal intestine were not preserved from fish in this study. This would have been useful for histological analysis of any gluten-induced enteritis.

Genetic predisposition is the primary risk factor for CD in humans, but there is evidence that early and abrupt introduction of gluten-containing foods into the infant diet alters the microbiome and increases likelihood of developing the disease (65). Several microorganisms are known to uptake taurine via Tau transporters for utilization as sulfur or carbon sources (66). Though no characterization has been performed of the cobia microbiome, such analysis has been performed in cats, another strict carnivore. Dietary studies have revealed that cats, like cobia, require dietary intake of taurine due to insufficient synthesis. By virtue of the fact that they have similar dietary needs (high protein: carbohydrate ratio) and metabolism (can only conjugate bile acids to taurine), there may also be common commensal bacteria. Similar to cats, alterations to populations of intestinal bacteria in cobia caused by intake of fermentable feed ingredients might increase deconjugation of bile salts and contribute to taurine depletion (67). Rapid shifts in the microbiome induced by diet are not isolated to carnivores, as it has also been well documented in humans along with celiac disease-associated dysbiosis (68, 69). In the case of juvenile cobia in Trial 1, it is possible that gut immaturity and incomplete establishment of commensal microbiota may have increased injurious effects from gluten. However, alterations to the normal carnivore microbial landscape from ingredients such as gluten could have the potential to affect the overall health of cobia at any stage of development.

The most significant physiological impacts of 4% dietary wheat gluten in European sea bass are the substantial increase in plasma taurine and induction of greater diversity of predominant taxonomic orders of intestine microbiota. These changes in no way suggest a poorer fitness outcome, and there were no significant differences in growth between the two groups. Therefore, there is no reason to contraindicate the addition of 4% wheat gluten addition to a completely plant-based diet for European sea bass.

The common plasma measures of overall health that differed between the 0 and 4% groups were calcium and AST. Plasma calcium was higher in the group fed 4% wheat gluten, whereas plasma AST levels were lower than they were in the fish fed the diet without wheat gluten. European sea bass stressed by hydrogen peroxide exposure exhibited higher levels of plasma calcium, so it could be indicative of a health effect (70). AST is an indicator of liver health and function, along with ALP (alkaline phosphatase) (71). In a fasting study in European sea bass performed by Peres et al., AST levels increased during the fasting period (36).

Our values for all plasma markers measured are similar to those reported by Peres et al. in European sea bass maintained on a fishmeal diet with the exception of cholesterol, ALP, and AST. For ALP and AST, our values were substantially lower, almost half of the values obtained by their group. This may be a function of a plant-based vs. fishmeal-based diet. Changes to cholesterol were not apparent in our study, but they have been in other studies with wheat gluten as a feed component. Plasma cholesterol decreased in European sea bass fed diets with graded levels of wheat gluten partially replacing up to 70% of the fishmeal as a protein source, but that may be due to the greater proportion of plant protein incorporation rather than an effect unique to wheat gluten (72). Interestingly, a partial replacement of fishmeal with corn as a protein source and no added wheat gluten was shown to raise plasma cholesterol and phospholipid levels. The authors attributed this effect to the higher carbohydrate content in the diet containing corn (73). Different plant sources of protein may have a variety of influences on plasma parameters.

The only change in measured tissue weights between diets was for the mid-intestine. There have been no reports of major histological changes to only the mid-intestine prompted by dietary ingredients. However, a gene expression study performed in European sea bass suggested that there is functional specialization across the length of the intestinal tract (47). Another study found that mid-intestine lactic acid bacteria (order *Lactobacillales*) are highly modulated by diets in a recirculating aquaculture system (74).

There are no studies correlating wheat gluten to higher levels of taurine in plasma. Recently there have been reports of diets for pets containing legumes marketed as “grain-free” causing cardiomyopathies related to taurine deficiency (75). It is possible that a feed ingredient such as wheat gluten might be influencing sequestration of taurine in plasma. Alternatively, endogenous levels of taurine may be increased as a result of dietary wheat gluten, perhaps to counter some pro-inflammatory effect of the wheat gluten.

The lack of detectable induction of higher levels of plasma IgM, and IgT suggest a lack of adaptive immune response. We also attempted to detect TNF-α in the plasma samples but obtained a high standard error suggesting that our antibody was non-specific. There was no detectable plasma factor capable of binding to gliadin. This is in contrast to our data presented in Chapter 2 for cobia consuming a diet containing less than 4% wheat gluten in which a gliadin-binding plasma factor(s) was produced.

The dietary inclusion of wheat gluten induced marked changes to the intestinal microbiome. In the intestines of fish fed 4% wheat gluten, there were increased predominant diversities of Proteobacteria as compared to the 0 wheat gluten group as well as the presence of Bacteroidetes in the mid- and posterior intestines. Several studies have linked dietary changes to alterations in the intestinal microbiome, and a small number have probed for this effect specifically with wheat gluten. Human studies in which participants shifted to a gluten-free diet showed variations to the microbiome, though the most significant variation was inter-patient. One of these studies found a significant decrease in the family *Veillonellaceae* of the *class Clostridia* (76)^-^(77). One study in zebrafish in demonstrated that fish fed diets containing wheat gluten (∼50% of formulation) had heightened abundances of *Legionellales, Rhizobiaceae,* and *Rhodobacter* over fishmeal-fed fish (78). In another study in zebrafish, fish fed wheat gluten had decreased *Bifidobacterium* relative to fish fed brine shrimp (79). Only an abstract could be located for this study, so the actual percentage of wheat gluten is unknown. In a study of Atlantic salmon, wheat gluten (∼14-20% of formulation) mixed with a legume protein (soybean meal or guar meal) increased abundance of lactic acid bacteria in the gut as compared to the reference fishmeal diet (80). Our data do not appear to correlate with these other studies, but many of the differences are likely attributable to the difference between species and the overall constitution of the diet: Plant-based, fishmeal-based, or a combination. Overall, the study results do not seem to indicate some type of disease state induced by the addition of 4% wheat gluten, as the fish have overall health comparable to that of the fish consuming a dietary containing no wheat gluten.

Taken together, these studies suggest that different species and life stages, even among the marine carnivores, may have different reactions to dietary ingredients such as wheat gluten. It is possible that the ability to synthesize taurine is significant in determining tolerance. Preliminary data from another study performed by our laboratory in European sea bass using diets containing 0 or 5% taurine suggest that unlike cobia, European sea bass are adequate endogenous synthesizers of taurine and do not require supplementation. Future studies of the effects of dietary wheat gluten in other marine carnivorous species with various taurine requirements will clarify this connection. Additionally, the observed partial amelioration of the negative effects induced by dietary wheat gluten in cobia should spur future research into taurine’s ability to mitigate effects elicited by other general stress factors.

## Acknowledgements

The authors would like thank the staff of the Aquaculture Research Center at the Institute of Marine and Environmental Technology: Steve Rodgers, Chris Tollini, and Joy Harris. We also thank Renate Reimschuessel, VMD, PhD, of the Food and Drug Administration for assistance with microscopy. This is contribution #XXXX from UMCES and #XX-XXx from IMET.

